# A history-dependent integrase recorder of plant gene expression with single cell resolution

**DOI:** 10.1101/2024.06.04.597320

**Authors:** Cassandra J. Maranas, Wesley George, Sarah K. Scallon, Sydney VanGilder, Jennifer L. Nemhauser, Sarah Guiziou

## Abstract

During development, most cells experience a progressive restriction of fate that ultimately results in a fully differentiated mature state. Understanding more about the gene expression patterns that underlie developmental programs can inform engineering efforts for new or optimized forms. Here, we present a four-state integrase-based recorder of gene expression history and demonstrate its use in tracking gene expression events in Arabidopsis thaliana in two developmental contexts: lateral root initiation and stomatal differentiation. The recorder uses two serine integrases to mediate sequential DNA recombination events, resulting in step-wise, history-dependent switching between expression of fluorescent reporters. By using promoters that express at different times along each of the two differentiation pathways to drive integrase expression, we successfully tied fluorescent status to an ordered progression of gene expression along the developmental trajectory. In one snapshot of a mature tissue, our recorder was able to reveal past gene expression with single cell resolution. In this way, we were able to capture heterogeneity in stomatal development, confirming the existence of two alternate paths of differentiation.

## Introduction

The sequential expression of genes during development progressively restricts and specifies a cell’s fate. Many of the genes involved in cell differentiation processes have been identified across a host of model and non-model organisms, and, in many cases, these genes have been assembled into cell fate pathways. However, a full understanding of development will require combining the establishment of these genetic regulatory networks with information about the extent and effect of variation in how these networks are experienced by individual cells. The key limiting technology to achieve this aim is a method to monitor gene expression history with single cell resolution that maintains spatial information.

Existing single cell RNA sequencing (scRNAseq) technologies provide a wealth of gene expression data in a variety of contexts^1^. However, scRNAseq provides only a snapshot of the cell’s transcriptional state at one point in time, fails to capture expression of lowly expressed genes, and requires destruction of the sample, thus losing spatial resolution. Recent methods combining scRNAseq with additional techniques (such as metabolically labeling mRNAs^2,3^; evaluating mRNA splice status^4^; and pseudo-time analysis^5^) have been developed to allow users to infer some temporal information and have been applied to study cell specification in *Zea mays*^6^ and *Arabidopsis thaliana*^7^. As a different approach to gain more information from scRNAseq, efforts to add spatial information include positional barcoding in tandem with next-generation sequencing in mammalian^8^ and plant^9^ systems; and *in situ* sequencing combined with *in situ* fluorescent hybridization in mammals^10^ and plants^11^. Such approaches provide some spatial context, but no temporal information and are quite costly and complex. Alternatively, keeping spatial and temporal information, methods which utilize fluorescent expression as a marker of genetic events have been widely used for decades in a diverse array of organisms including mouse models^12^ and plants^13^. These approaches are limited in information due to: the finite number of fluorescent tags that can be used at one time; the stress to the organism due to repeated irradiation; and the long half-lives, slow maturation time, and photobleaching risk of fluorescent proteins. One study^14^ successfully implemented a fluorescence-based live imaging approach over multiple days for studying lateral root development in *Arabidopsis,* but was able to track only six individuals due to the complexity of the imaging setup.

Recently, DNA-based recording systems have been developed and overcome some of the challenges of ‘omic and microscopy-based technologies. Systems based on CRISPR-Cas9 mediated mutations and subsequent DNA sequencing enabled recording of the Wnt signaling pathway in mammalian cells^15^ although this approach lacked spatial resolution. DNA-based recording devices utilizing CRISPR have also been used for lineage tracing, for example in reconstructing lineages in *Arabidopsis* tissues^16^, zebrafish organs^17^, and in development of a lineage tracing mouse line^18^. Researchers have also utilized the DNA recombination abilities of integrases for similar DNA-based recording approaches. For example, the Cre-LoxP system has been employed to trace phloem and xylem vascular cell lineage in *Arabidopsis*^19^. Other work employs stochastic integrase recombination to generate barcodes which are then deconvoluted to construct cell lineages in mouse cell lines and *Drosophila melanogaster*^20^. Synthetic memory switches using tissue specific integrase expression have recently been implemented in *Arabidopsis*^21,22^. More complex integrase circuits have been designed and implemented in bacterial cells^23–25^, mammalian cells^26,27^, and plant protoplasts^21,27^.

Plants are particularly interesting organisms for studying the differentiation process as they continue to generate new organs throughout their lifetime. Decoding plant developmental programs can also inform engineering of novel crops with enhanced properties to adapt to environmental stresses accelerated by climate change^28^. Here, we built on previous work using single serine integrase switches prototyped in *Nicotiana benthamiana*^29^ and implemented in *Arabidopsis* for tracking the expression of genes underlying lateral root initiation^22^. Following the logic of a previously engineered recorder in bacterial cells^25^, we engineered a four-state history-dependent integrase-based recorder of gene expression in *Arabidopsis*. We first validated the design of the recorder target using constitutive expression of two serine integrases: PhiC31 and Bxb1. We then tested the efficacy of the circuit to record two well-known transcriptional trajectories, those occurring during the development of lateral roots and stomata. In both cases, we were able to successfully capture the order of expression of two genes in individual cells. Additionally, the recorder revealed heterogeneity in the gene expression history of mature stomata, highlighting the alternative differentiation pathways that they had experienced.

## Results

### An integrase-based history-dependent recorder functions in plants

Serine integrases mediate site specific DNA inversions or excisions based on the presence and orientation of specific integrase sites (inversion when sites are in opposite orientation and excision when in the same orientation). Our integrase-based, history-dependent recorder of gene expression leverages these recombination abilities, with four possible fluorescent expression outputs, each indicative of a specific order of occurrence of signal inputs. The recorder is composed of two parts: a target (composed of integrase sites, fluorescent proteins, and gene regulatory elements) and an integrase construct (mediating integrase expression). We designed our target construct (Fig. 1a) for *Arabidopsis* with a strong constitutive p35S promoter^30^ and three genes encoding fluorescent proteins: ER-localized mtagBFP2^31^ (named BFP from here on), nuclear-localized mCherry^32^ (named RFP from here on), and nuclear-localized NeonGreen^33^ (named GFP from here on). The promoter and reporter genes are flanked by integrase sites for the orthogonal serine integrases: PhiC31 (Fig. 1a, gray triangles) and Bxb1 (Fig. 1a, black triangles). The target has four possible DNA states which result in different expression outputs: State 0 (BFP expression), State 1 (RFP expression), State 2 (GFP expression), and State α (no fluorescent expression). The switches between states are order-dependent and mediated by the PhiC31 and Bxb1 integrases.

**Figure 1:**
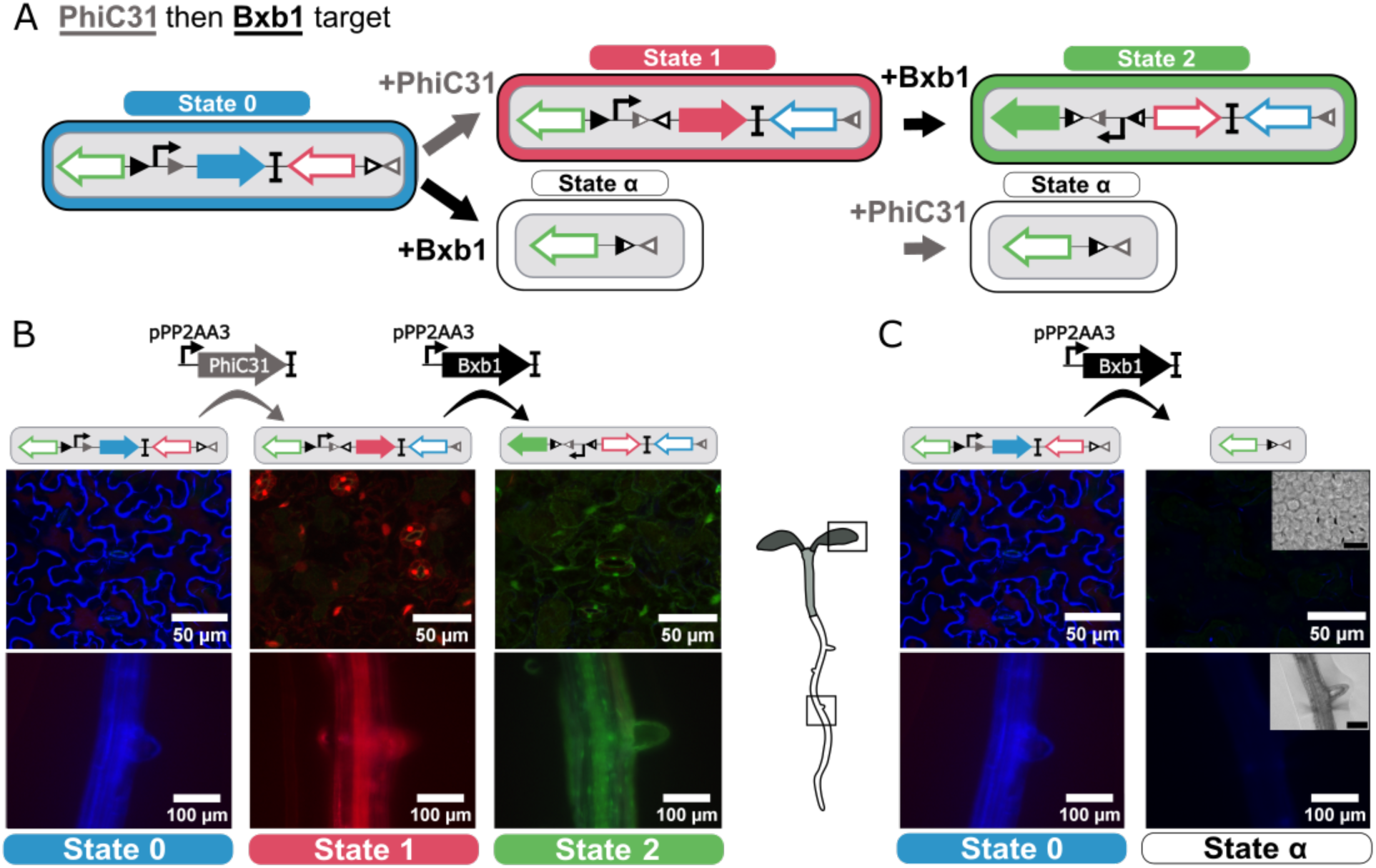
The history dependent integrase circuit switches between fluorescent states based on order of inputs. **A** Schematic of integrase target circuit. The target consists of: two sets of integrase sites for the PhiC31 and Bxb1 integrases; three genes for fluorescent proteins (mtagBFP2 (BFP), mCherry (RFP), and NeonGreen (GFP)); and a p35S strong constitutive promoter for fluorescent expression. The initial state of the target (State 0) results in expression of mtagBFP2. Upon expression of the PhiC31 integrase, an inversion causes the target to switch to State 1 and results in expression of RFP (State 1). Subsequent Bxb1 expression causes another inversion, flipping the promoter direction and mediating the switch to State 2 which results in GFP expression (State 2). Alternatively, addition of the Bxb1 integrase in a cell with the State 0 target configuration results in a DNA excision, leading to a switch to State α and a loss of fluorescence. **B** The PhiC31-Bxb1 integrase order mediates a series of switches from State 0 to State 1 to State 2. (left) The initial target in State 0 shows strong BFP expression in the leaf and the root. (middle) Constitutive expression of PhiC31 mediates a switch of the target to State 1 and strong RFP fluorescence in leaf and root tissue. (right) Subsequent constitutive expression of Bxb1 mediates a switch from State 1 to 2 and results in strong GFP expression in leaf and root tissue. **C** Expression of Bxb1 prior to PhiC31 mediates an “out of order” switch to State α and loss of fluorescence. (left) The initial target in State 0 shows strong BFP expression in the leaf and the root. (right) Constitutive expression of Bxb1 with the target in State 0 results in a switch to State α and a loss in fluorescence in the leaf and root. Black scale bars in the brightfield images correspond to 100 μm (root) and 50 μm (leaf).

The PhiC31 then Bxb1 target allows tracking of the PhiC31 then Bxb1 lineage, switching progressively from State 0 to State 2 when PhiC31 is expressed before Bxb1. In this target, State 0 represents the initial DNA state with no integrase expression. Expression of PhiC31 integrase mediates the switch to State 1 via inversion of the BFP-RFP cassette, resulting in a switch from BFP to RFP expression. Subsequent expression of Bxb1 integrase in these State 1 cells mediates the subsequent switch from State 1 to 2 via inversion of the p35S promoter and results in GFP expression. If Bxb1 is expressed “out of order” prior to PhiC31 in a cell, it excises 75% of the target, including the p35S promoter, corresponding to switch from state 0 to state α and resulting in a loss of fluorescence (Fig. 1a). The integrase switches are irreversible and heritable, so descendent cells inherit the target DNA state from the mother cell. To characterize this target, we used a constitutive promoter to drive each integrase separately, the promoter for PROTEIN PHOSPHATASE 2 A SUBUNIT 3 (pPP2AA3, commonly used as a control for qPCR^34^). pPP2AA3 is not as strongly expressed as other commonly used plant constitutive promoters such as pUBQ10 or p35S, making it a better match for the expression level of most developmental genes.

We first transformed the target in *Arabidopsis*, to obtain a stable target line. We confirmed strong expression of only BFP in both root and leaf tissue (Fig. 1b, left; Fig. 1c, left) in the target plant line selected for the rest of the experiments. We subsequently transformed this plant line with a PhiC31 constitutively expressed construct. We observed a switch in expression to RFP in both root and leaf tissue (Fig. 1b, middle) as expected. To test the State 1 to State 2 switch in isolation, we designed a preswitched target construct where the starting state of the target is the DNA State 1. We successfully confirmed that this preswitched target shows strong RFP expression in the leaf and root (Fig. S1). When transformed with a pPP2AA3-expressed Bxb1 construct, this preswitched target exhibited strong constitutive GFP expression in both the leaf and root (Fig. 1b, right) confirming the switch from State 1 to State 2. Transformation of the State 0 target with pPP2AA3-expressed Bxb1 resulted in a strong reduction in BFP expression in the T1 generation (Fig. 1c) as expected. To confirm the switch from State 0 to State α, we genotyped this seedling and confirmed the presence of the target in a DNA state corresponding to the length of the State α target (Fig. S2).

In addition to the PhiC31 then Bxb1 target, we also generated a Bxb1 then PhiC31 target (Fig. S3) wherein the Bxb1 then PhiC31 lineage is tracked, switching from State 0 to 1 then 2 when Bxb1 then PhiC31 are expressed sequentially. We validated the initial switches for this target, showing Bxb1 expression causes a switch to RFP expression, therefore from State 0 to State 1 in the target, and PhiC31 expression to no fluorescent expression, therefore to State α of the target (Fig. S3). We also developed and made publicly available an alternate target for tracking the PhiC31 then Bxb1 lineage in which the reporters are expressed with pUBQ10 instead of p35S.

### History dependent recording of gene expression during lateral root initiation

For the next step in prototyping, we used well-characterized developmental promoters in the integrase construct to control expression of the integrases. For this test, we turned to the development of lateral roots, as many promoters for genes involved in lateral root cell fate specification have been already characterized using reporter constructs^35,36^ and, for a few promoters, using integrase switches^22^. Beyond the depth of knowledge about lateral root developmental programming, it is also of high interest for engineering efforts aimed at increasing drought resilience^37^.

Lateral roots develop from a small subset of xylem pole pericycle (XPP) cells, designated as “founder cells”. These founder cells then undergo a series of proliferative, asymmetrical cell divisions, establishing the new lateral root^38^. Our lateral root recorder was designed to track the expression of two genes: *ARABIDOPSIS HISTIDINE PHOSPHOTRANSFER PROTEIN 6* (*AHP6*) and *GATA TRANSCRIPTION FACTOR 23* (*GATA23*). AHP6 is a negative regulator of cytokinin signaling, and it plays an important role in guiding protoxylem formation^39^ and orienting cell divisions during lateral root initiation. It is expressed in the xylem poles and XPP cells^40^. *GATA23* is strongly expressed in lateral root founder cells, setting into motion the series of cell divisions which form the lateral root^41^. For the integrase construct component of the lateral root recorder, we placed the first integrase (PhiC31) under control of pAHP6 and the second integrase (Bxb1) under control of pGATA23.

First, we characterized the single switches for pAHP6 and pGATA23. To characterize the performance of the single switches, we categorized seedlings as: (1) ‘as expected’ meaning the only RFP expressing cells were those which expressed the recorded gene (or descended from such a cell); (2) overswitched meaning additional cells were expressing RFP; and (3) underswitched meaning that less cells were expressing RFP (see Table S1 for an overview of these categories for each single integrase switch and full recorder). For our single switches and recorder lines, we characterized all T2 seedlings without selection for integrase constructs or homozygosity. Therefore, we observed unswitched T2 seedlings with all cells expressing BFP. We attributed this to loss of the integrase construct between generations, as the proportion of these unswitched seedlings is consistent with that for prior integrase switches^22^ and is consistent with Mendelian heritability (around 25%)^42^, so we omitted these seedlings from our characterization (see Source Data file 1).

For pAHP6, we transformed the PhiC31 then Bxb1 target line with a single integrase (PhiC31) under the control of the pAHP6 promoter, which we called the pAHP6 single integrase switch, as it only uses the switch from State 0 to State 1 of the target (Fig. 2a). In any cell where *AHP6* has been expressed, that is, cells in the xylem poles and XPP, we expected to see a switch from BFP to RFP expression, indicating a switch of the target from State 0 to State 1. Because the switch is heritable, we would predict that in a plant with this single pAHP6 switch, all xylem pole cells, XPP cells and their descendants (e.g., lateral roots) would express RFP, representing the ‘as expected’ switch pattern (Table S1). Because no switch was observed initially in the xylem poles or XPP in any T1 seedlings, we added an NLS (nuclear localization signal tag^22^) onto PhiC31 to increase its DNA recombination activity, resulting in four out of seven T1 seedlings with the ‘as expected’ switch pattern (Table S2). Using this integrase construct with the NLS, we were able to consistently generate ‘as expected’ T2 seedlings (Fig. 2a). In the best performing pAHP6 single integrase switch T1 line (with NLS), 78% of T2 seedlings showed the expected switch pattern and the worst performing line had 18% with the expected pattern (Fig. 2a, Fig. S4a). Part of this seedling to seedling variability is due to the fact that we characterized a mix of hetero and homozygous seedlings for the integrase construct. There were also seedlings consistently in the underswitched category (the line with the highest prevalence had 59% underswitched), in this case meaning the switch occurred in the lateral root but not in the xylem poles or XPP cells. This indicates stronger expression of *AHP6* in lateral root precursor cells, consistent with the fact that *AHP6* expression is induced by the plant hormone auxin^40^ which is found at high levels in founder cells during early lateral root initiation^43^.

**Figure 2:**
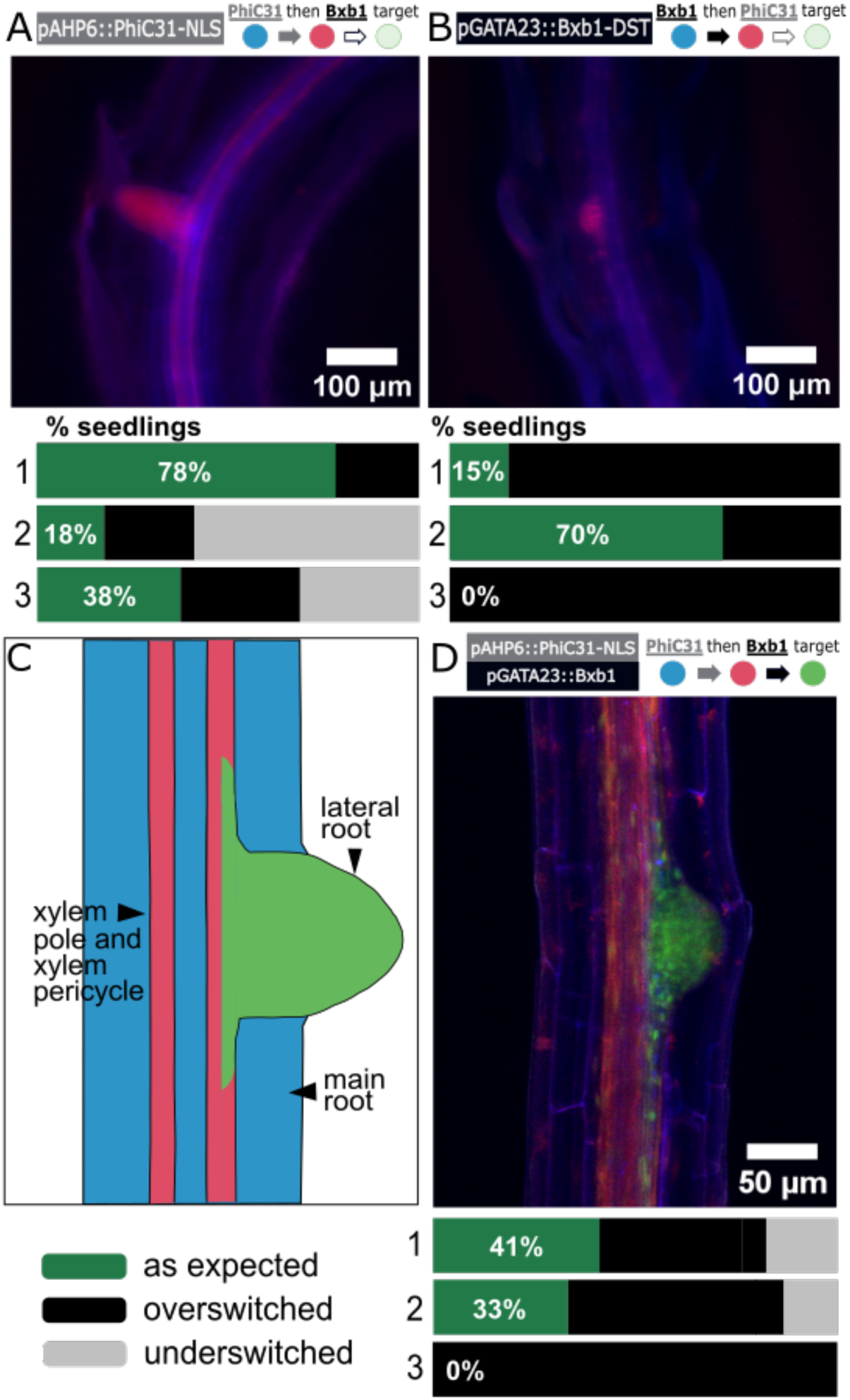
Application of the history-dependent recorder in lateral roots. **A** A pAHP6 single integrase switch. Plants carrying the PhiC31 then Bxb1 target were transformed with pAHP6::PhiC31-NLS. A representative image of a T2 seedling is shown with the switch characterization for three representative lines (line 1: T2P5 (n=17), 2: T2P2 (n=16), 3: T2P7 (n=9)), see Fig. S4a), where n represents the number of seedlings characterized in each line, omitting seedlings with no switching. The bar graphs represent the proportion of seedlings in each T1 line showing the ‘as expected’, overswitched, and underswitched switch pattern (as defined in Table S1). **B** A pGATA23 single integrase switch. Plants carrying the Bxb1 then PhiC31 target (Fig. S3) were transformed with pGATA23::Bxb1-DST. A representative image of a T2 seedling is shown with the switch characterization for three representative lines (line 1: T2P5 (n=17), 2: T2P3 (n=20), 3:T2P1 (n=16), see Fig. S4a). The bar graphs represent the proportion of seedlings in each T1 line showing the ‘as expected’, overswitched, and underswitched switch pattern (as defined in Table S1). **C** Predicted recorder output. Xylem pole and xylem pole pericycle cells should express RFP (State 1, red), the cells in the lateral root should express GFP (State 2, green), and any other cell should remain in the initial BFP-expressing (State 0, blue). **D** Lateral root recorder output. T2 seedlings from 5 lines which came from T1 plants with an ‘as expected’ or underswitched recorder output were screened. A representative image for one of these T2 seedlings is shown as well as characterization of the recorder output for three representative T1 lines (line 1: T2P1 (n=17), 2: T2P3 (n=15), 3: T2P9 (n=13), see Fig. S5a). The bar graphs represent the proportion of seedlings in each T1 line showing the ‘as expected’, overswitched, and underswitched switch pattern (as defined in table S1). Source data is provided as a Source Data file 1. Additional microscope images can be found in Source data file 2.

*GATA23* has been previously used to drive a single PhiC31 integrase switch as a recorder of lateral root initiation^22^. To characterize a Bxb1-mediated single switch, we transformed the Bxb1 then PhiC31 target (Fig. S3) with a single Bxb1 construct driven by pGATA23 such that Bxb1 mediates the switch from State 0 to 1 (Fig. 2b). As only one switch of the target is utilized, we refer to this as the pGATA23 single integrase switch. Consistent with our knowledge of *GATA23* expression and previously characterized *GATA23* integrase switch^22^, we expected to observe RFP expression only in cells within the lateral root, constituting the ‘as expected’ switch category (Fig. 2b, Table S1). The pGATA23::Bxb1 single integrase switch resulted in T1 seedlings with RFP expression in cells outside the lateral root (overswitched) (Table S2). To achieve switch specificity consistently in the T1 and T2 generations, we added an RNA destabilization tag from SMALL AUXIN UP-REGULATED RNA genes^44^ (DST) to the Bxb1 construct^22^. This pGATA23::Bxb1-DST single integrase switch resulted in five out of eight T1 seedlings with the ‘as expected’ switch pattern (Table S2). From these ‘as expected’ T1s, we obtained three T1 lines out of five with seedlings behaving ‘as expected’ (ranging from 15% to 71% of seedlings) (Fig. 2b, Fig. S4). The majority of the seedlings in most of the lines were overswitched. The T1 line with the lowest proportion of overswitching had 29% and the highest line was 100% overswitched. The full T2 characterization of the single switches for the lateral root genes can be found in Fig. S4a.

Next, we generated the full lateral root recorder, building the dual integrase construct with pAHP6-driven PhiC31 and pGATA23-driven Bxb1 and transforming it into the PhiC31 then Bxb1 target line. We added an NLS to the pAHP6::PhiC31 construct and no tag for pGATA23::Bxb1 construct. According to the known expression patterns of *AHP6* and *GATA23*, we expected to observe xylem pole and XPP cells expressing RFP, lateral root cells expressing GFP, and the rest of the cells in the root should remain in their initial, BFP-expressing state (Fig. 2c). We were able to consistently generate seedlings with the ‘as expected’ pattern (Fig. 2d). In the T1 generation (Table S3), four out of nine seedlings were overswitched with the rest being some combination of ‘as expected’ and underswitched. In the context of the full lateral root recorder, overswitched refers to seedlings with PhiC31 and/or Bxb1 overswitching, with GFP-expressing cells outside the lateral root and/or RFP-expressing cells outside the xylem poles and XPP (Fig. S6).

We characterized T1 lines from the ‘as expected’ and underswitched T1s because subsequent generations are likely to have stronger integrase expression and, as a result, we would expect an increase in overswitching between generations. In the T2 generation, an average of 23% of seedlings screened per line showed the expected output. Those that deviated from the expected phenotype consisted primarily of overswitched seedlings (Fig. S5a). In contrast to the pGATA23 single switch where the pGATA23::Bxb1 construct led to overswitching in every T1 seedling (Table S2), in the full recorder we were able to achieve switching specific to *GATA23* expression without addition of a DST to Bxb1. In our dual integrase construct the Bxb1 transcriptional unit is immediately downstream of the PhiC31 unit, and this construct architecture has been shown to result in reduced expression of the downstream gene^45^ possibly due to transcription-induced DNA supercoiling^46^. Schematics of all the observed switch patterns can be seen in Fig S6. Overall, we successfully implemented the recorder in tracking the expression patterns of *AHP6* and *GATA23* during lateral root initiation and were able to consistently generate T2 seedlings with the expected switch pattern.

### History dependent tracking of gene expression during stomatal development

We next applied our history-dependent recorder to a different well-characterized differentiation program: stomatal development. Stomata are each composed of two guard cells which together form a small pore in the leaf surface. These pores are required for efficient gas exchange and are a target of interest for engineering increased carbon capture with higher water use efficiency^47^.

Each stoma descends from a proliferating meristemoid cell on the leaf epidermis which serves as the precursor of the stomatal cell lineage. Stomata develop following a stereotypical progression wherein the meristemoid undergoes some number of asymmetrical cell divisions before dividing symmetrically exactly once into the two guard cells that comprise the stoma. Many of the genes involved in this progression from meristemoid to guard cell have been extensively studied^48^. We focused on recording the expression of two genes in the context of stomatal development: *SPEECHLESS* (*SPCH*), which enables the asymmetrical cell divisions of the meristemoid, and *MUTE*, which is expressed later and is essential to trigger the single symmetric division into two guard cells^49^. We therefore used pSPCH to drive expression of the first integrase (PhiC31) and pMUTE to drive expression of the second integrase (Bxb1). The final recorder line contains this integrase construct and the PhiC31 then Bxb1 target.

We first separately tested the individual integrase switches with pSPCH driving PhiC31 and pMUTE driving Bxb1. For the pSPCH single switch, we transformed the pSPCH-driven PhiC31 construct into the PhiC31 then Bxb1 target line, expecting to observe a switch from BFP to RFP expression (State 0 to 1) in the guard cells plus any surrounding epidermal cells that resulted from asymmetric meristemoid divisions (the ‘as expected’ switch category) (Fig. 3a, Table S1). In this pSPCH single integrase switch, overswitched means that additional leaf epidermal cells were expressing RFP, and underswitched means that less cells were expressing RFP (Table S1). In the T1 generation, four out of the ten seedlings showed ‘as expected’ switching (Table S4). In the T2 generation, we obtained up to 50% of seedlings per line switching ‘as expected’ with a high variability between lines. Most of the T2 seedlings showed an overswitched phenotype, with some or all leaf epidermal cells not in contact with stomata expressing RFP. The most overswitched T1 line had 100% overswitched seedlings while the lowest rate of overswitching observed was 14%. One line had 64% of the seedlings underswitched, with only the guard cells of the stomata expressing RFP (Fig. 3a, Fig. S4b).

**Figure 3:**
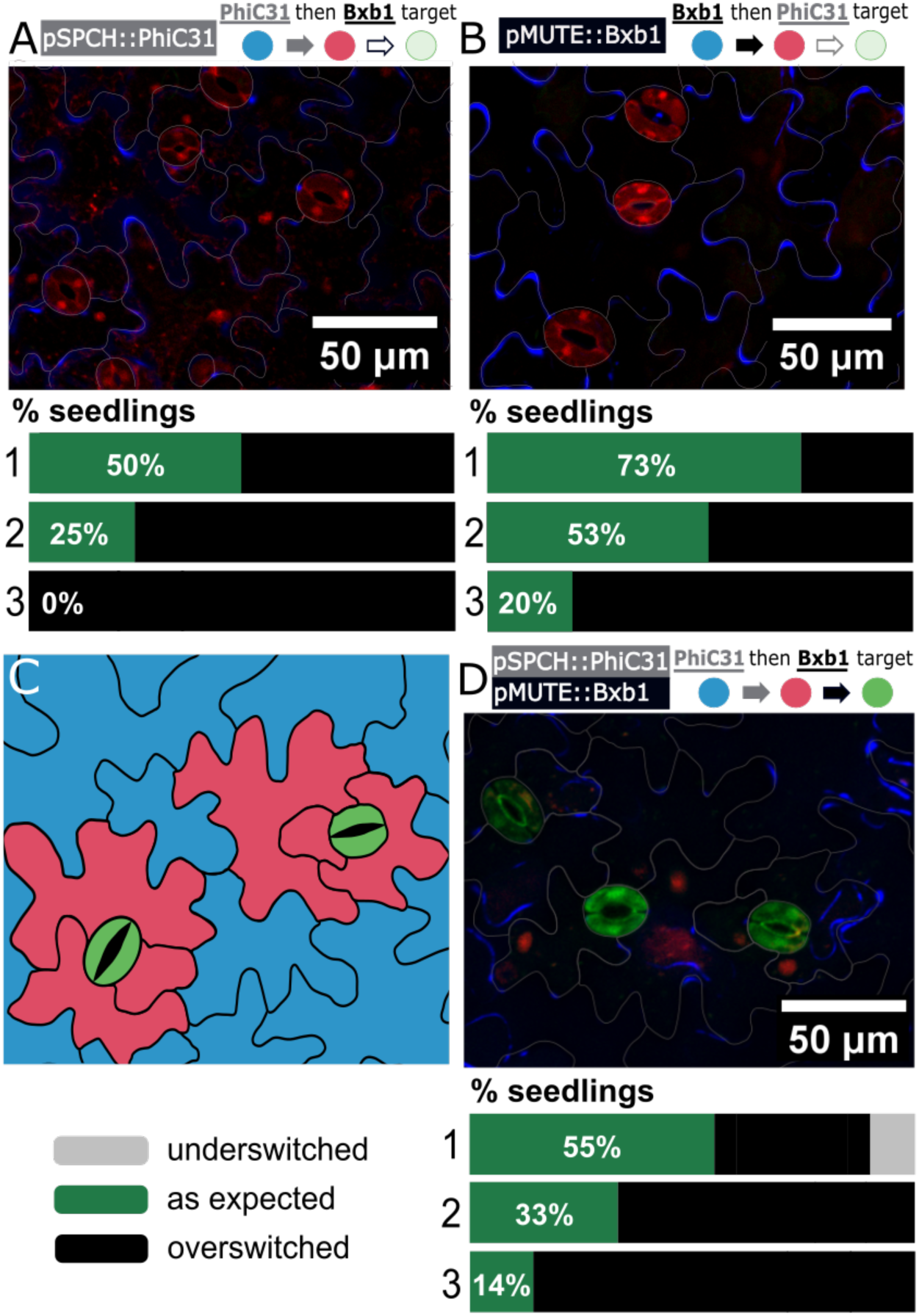
Application of the history-dependent recorder during stomata development. **A** A pSPCH single integrase switch. Plants carrying the PhiC31 then Bxb1 target were transformed with pSPCH::PhiC31. A representative image of a T2 seedling is shown as well as the switch characterization for three representative lines (line 1: T2P3 (n=12), 2: T2P8 (n=14), 3: T2P1 (n=14), see Fig. S4b) where n represents the number of seedlings characterized in each line omitting seedlings with no switching. **B** A pMUTE single integrase switch. Plants carrying the target were transformed with pMUTE::Bxb1. A representative image of a T2 seedling is shown with the switch characterization for three representative lines (line 1: T2P2 (n=15), 2: T2P7 (n=15), 3: T2P1 (n=15), see Fig. S4b). **C** Prediction of stomatal recorder output. Guard cells are predicted to be in State 2 and expressing GFP (State 2, green), surrounding epidermal cells that are the result of meristemoid division should be expressing RFP (State 1, red), and any other cell should sustain the initial BFP expression (State 0, blue) . **D** Experimental stomatal development recorder output. The PhiC31 then Bxb1 target is transformed with the dual stomatal integrase construct. One representative image of a T2 seedling matching the expected output is shown. The recorder performance is shown for three representative lines (line 1: T2P1 (n=20), 2: T2P4 (n=18), 3: T2P2 (n=20), see Fig. S5b), with percentages indicating the proportion of seedlings matching the expected recorder output and n representing the number of seedlings characterized in each T1 line omitting seedlings with no switching. Source data are provided as a Source Data file 1. Additional microscope images can be found in Source data file 2.

To generate the pMUTE single switch, we built a pMUTE-driven Bxb1 construct and transformed it into the Bxb1 then PhiC31 target (Fig. S3) line. We characterized the lines similarly as for the pSPCH single switch lines in the T1 and T2 generations with the ‘as expected’ phenotype being RFP expression in the guard cells of the stomata only (Table S1). In the T1 generation, five out of ten seedlings showed the ‘as expected’ switch pattern (Table S4). In the T2 generation, we consistently obtained a majority of seedlings expressing RFP only in the guard cells (73% and 53%) (Fig. 3b). Every seedling which did not fit this pattern showed an overswitched phenotype, with RFP expression in additional leaf epidermal cells (Fig. 3b, Fig. S4b).

We then generated the full stomatal recorder by building the dual integrase construct with pSPCH-driven PhiC31 and pMUTE-driven Bxb1 and transforming it into the PhiC31 then Bxb1 target line. We expected to see expression of GFP in the guard cells of the stomata, expression of RFP in any surrounding epidermal cells that resulted from asymmetric meristemoid divisions, and expression of BFP in the remaining leaf epidermal cells which are not involved in stomatal differentiation. Of the nine T1 seedlings generated, six showed GFP expression specific to stomata and the lines from these six seedlings were characterized (Table S3). In five out of the six lines, we obtained seedlings with the expected phenotype (Fig. 3d, Fig. S5b), with a proportion of up to 50%, and a low of 14% ‘as expected’ seedlings per line, with three out of six lines having an ‘as expected’ proportion greater than 33%. The majority of the not ‘as expected’ seedlings were overswitched, meaning they were expressing GFP in cells other than the guard cells and/or expressing RFP in excess epidermal cells. The overswitched category accounted for the majority of seedlings in most of the stomatal recorder T1 lines, including 100% of seedlings in one line, with a low of 35% and a median of 68% of seedlings. Most of the overswitched seedlings (median of 67%) showed both PhiC31 and Bxb1 overswitching (Fig. S7). A small proportion of seedlings (15% or less) showed PhiC31 underswitching, meaning the guard cells were expressing GFP and the rest of the epidermal cells were expressing BFP, but no cells were actively expressing RFP. For one line, we obtained only seedlings with non-specific switching (both PhiC31 and Bxb1 overswitching), meaning some or all non-stomata epidermal cells were expressing GFP (switched to State 2) and/or some or all epidermal cells not in contact with at least one stoma were expressing RFP (switched to DNA State 1). A schematic summary of all the observed recorder outputs is in Fig. S7. As highlighted previously, some of the variability between seedlings is due to differences in zygosity of the integrase construct. Regardless, we consistently generated seedlings in the ‘as expected’ category for both recorders.

### The history dependent recorder can identify cells that have undergone an alternate developmental path

For the recorder seedlings with an ‘as expected’ phenotype, the majority of the stomata expressed GFP as expected; however, there was a subset of stomata which did not express any fluorescent protein (Fig. 4a). In the design of the recorder, a cell which has no fluorescence indicates that the target has been switched to State α due to Bxb1 recombination before PhiC31 recombination. This result suggests the possibility that there are two stomatal populations which differ in how likely they are to switch to State α versus to proceed through the series of switches to State 2. We hypothesized that this discrepancy was due to differences in the relative timing of *SPCH* and *MUTE* expression. A closer relative timing of expression would increase the chances of Bxb1 mediating the switch to State α, while a longer timing would give PhiC31 more of a head start, increasing the likelihood of the cell to switch to State 1 and then State 2. Indeed, temporal logic gates based on similar integrase designs have been employed previously in *E. coli* to reveal the timing between events^50^.

**Figure 4:**
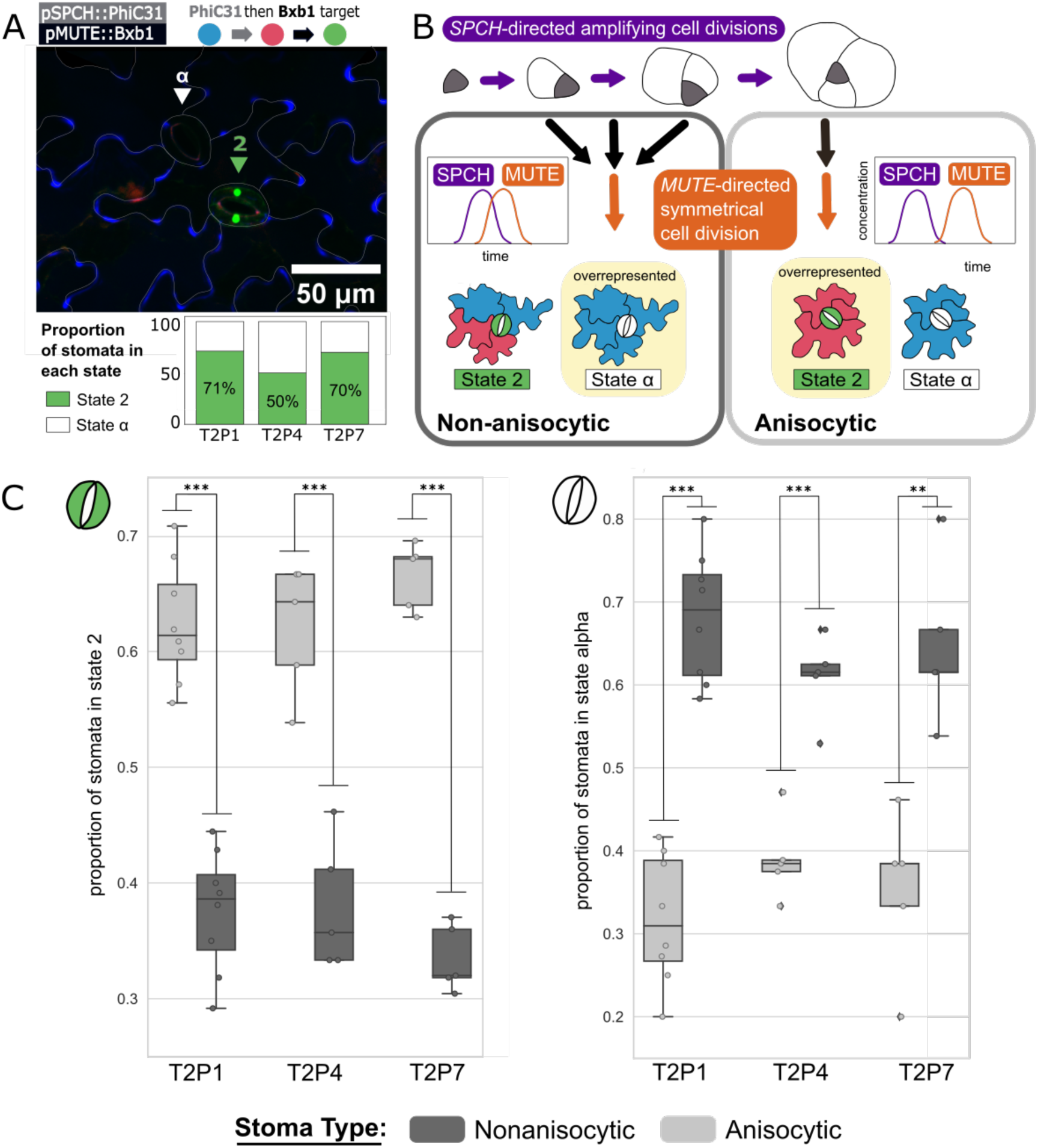
The history dependent recorder reveals two distinct stomatal populations. **A** State 2 and State α stomata. A confocal image shows two stomata: one in State 2 (green triangle marker) and one in State α (white triangle marker). Below the image, the total proportion of stomata in each state is shown in a barplot with each bar representing one of the three best performing stomata recorder lines (labeled on the x axis). The total number of stomata counted was 257, 172, and 174 respectively for the T2P1, T2P4, and T2P7 lines. **B** Alternate paths of stomatal development. The stereotypical understanding of stomatal development follows an anisocytic (A) scheme, with the meristemoid dividing asymmetrically three times before dividing symmetrically into two guard cells. Non-anisocytic (NA) stomata divide asymmetrically zero, one, or two times before the terminal symmetric division. Recorder output is predicted to differ between stomata type due to varied timing of integrase onset. An A stomata is surrounded by exactly three cells of unequal size and should show the recorder output as laid out in Figure 3. An NA stomata can have two, four, five or three equally sized neighbor cells and is expected to have the guard cells in state α. **C** Characterization of recorder state in A vs. NA stomata population. The boxes represent the proportion of each type of stomata (anisocytic in light grey and non-anisocytic in dark grey) within each of the three best performing stomata recorder lines that are in State 2 (left graph) and State α (right graph) for one line. Each point represents the proportion of stomata in each state for one T2 seedling. The x axis shows the three T1 lines that were characterized. For each seedling, a minimum of 15 of each stomata type was counted (at least 30 total stomata per seedling). 10 T2 seedlings were counted per line. To evaluate statistical significance between the result for A and NA stomata populations, a Student’s t test was performed (*p<0.05, **p<0.01, ***p<0.001). Source data are provided as a Source Data file 1.

It has been shown that the timing between *SPCH* and *MUTE* expression differs between the development of two types of stomata: anisocytic (A) and nonanisocytic (NA)^51^. During stomatal differentiation, *SPCH* initiates and *MUTE* terminates the asymmetrical divisions of the meristemoid^49^. Closer timing of expression between *SPCH* and *MUTE* permits fewer asymmetrical divisions, resulting in an NA stoma, whose development can include zero, one, or two asymmetric meristemoid divisions. In contrast, a longer delay between *SPCH* and *MUTE* expression allows the full three asymmetrical cell divisions needed for development of an A stoma (Fig. 4b). As a result, A stomata, the majority of stomata in most cases, are easily identified because the series of three asymmetrical divisions results in a stoma surrounded by exactly three cells of unequal size, whereas an NA stomata can have two, four, or five neighbor cells or three neighbor cells of roughly equal size^52^ (Fig. 4b).

We hypothesized these two developmental lineages, with their differences in relative gene expression timing, led to two stomata populations with different proportions of cells in each recorder state. We predicted that the State 2 stomata should be overrepresented in the A population and State α stomata should be overrepresented in the NA population. To test this hypothesis, we counted 25 stomata per T2 seedling in 10 individuals. This was repeated for the three best performing stomata recorder lines, only characterizing seedlings matching the “as expected” phenotype (Fig. S7). For each counted stoma, we categorized it as A or NA and State 2 or State α. We then calculated for both stomatal populations, the percentage of stomata in State 2 versus in State α. For A stomata, an average of 65% across all three lines were in State 2 and 35% in State α. In contrast, for NA stomata, an average of 33% across the three lines were in State 2, with 67% in State α (Fig. 4c). Therefore, as predicted, State 2 stomata were enriched in the A population compared to NA where a majority of stomata were in State α. In theory, the State α stomata could be a result of stomatal differentiation which skips *SPCH* expression entirely. If this were the case, the pSPCH single integrase switch result (Fig. 3a) would include unswitched stomata at a comparable rate, however we observed that all stomata in the ‘as expected’ pSPCH single integrase switch lines expressed RFP (see Source Data file 1) and therefore include *SPCH* expression in their lineage. This result supports our hypothesis that the disparity in recorder output was split probabilistically based on the extent of asymmetrical cell divisions prior to *MUTE* expression.

## Discussion

Developmental biology has made enormous strides in identifying gene regulatory networks that specify cell fate at a population scale (i.e., for the ‘average’ cell with a given identity). There is increasing interest in understanding the history of individual cells, including quantifying cell-to-cell variation in gene expression dynamics and context-specific, cell non-autonomous influences on cell fate. We have built a tool that can help with these efforts. Our integrase-based history-dependent recorders worked in two distinct developmental contexts: the initiation of a new root from the pericycle cells of the primary root and the differentiation of stomata in the leaf epidermis. In both cases, we could readily visualize cells in State 0, State 1 and State 2, and we consistently generated seedlings whose fluorescent expression matched our expectations based on known gene expression patterns. In tracking stomatal development, we also detected cells with no fluorescence, indicating a switch to State α. This finding supports prior evidence for the existence of two populations of stomata. Our results supported the hypothesis that the difference between stomata in State 2 and State α reflect differences in the relative dynamics of expression of the two genes used to make the recorder. Collectively, our work demonstrates the utility of the recorder for revealing gene expression variation even in seemingly uniform cell differentiation processes.

For a targeted study of the gene expression patterns underlying cell development paths of interest, our recorder provides an accessible and powerful methodology. Our recorder was sensitive enough to capture a disparity in gene expression timing in a very well-studied context, so it should prove even more beneficial for those working in less studied cell development contexts. This sensitivity combined with the strong constitutive, sustained expression of the reporters allows the recorder to act as an amplifier of the gene expression signal, which should prove useful for studying low expressing genes not suited for traditional transcriptional reporters, and enables evaluation of developmental events over timescales far exceeding the degradation time of the fluorescent proteins. Our recorder enables a snapshot readout of each cell’s gene expression history with only standard molecular biology techniques and access to a fluorescent microscope to take images at just a single or small number of time points.

While much information could be gleaned from the recorders analyzed here, future engineering efforts, especially those focusing on less well-characterized pathways, may require a systematic approach to reduce variability. This issue is a persistent one for engineering in plants. Line-to-line variation is likely due to the random insertion location of transgenes leading to different levels of gene expression. In the future, this issue could be addressed using targeted DNA transformation techniques. Variability in performance between seedlings carrying the same transgene insertions could be addressed with iterative rounds of optimization with our previously characterized tuning parts^22^. In this study, variability is also likely exacerbated by varied integrase expression levels among seedlings within T1 lines due to differences in zygosity of the integrase construct.

A four-state recorder like the designs used here can capture the history of expression for two genes. Using our current target design, researchers could generate multiple recorders within the same cell specification pathway to capture the expression patterns for more gene combinations. Moreover, the current recorder infrastructure could be adapted following existing integrase target designs to generate a target with a higher number of inputs^25^. The design capabilities of integrase circuits^23^ combined with the large number of characterized orthogonal integrases^27,53^ should enable construction of circuits with higher numbers of outputs. The possible scope is limited in principle only by the availability of spectrally distinct reporters, and this limit could be overcome with multiplexing techniques^54^ or switching to DNA barcodes^20^.

In theory, our recorder designs should be transferable to any plant that can be transformed, for applications such as those facilitating evo-devo studies of key developmental processes such as stomata.Our recorder can be applied in plants which undergo different paths of stomatal development, such as in amphibious plants which largely skip the *SPCH*-driven, asymmetrical cell divisions^55^ and grasses which have a completely different arrangement of subsidiary cells compared to *Arabidopsis*^56^. In addition, integrase switches can be adapted to allow study of essential genes in non-essential organs^57^. Porting this system into crop plants could enable optimization and quantification of traits associated with climate resilience, facilitating marker-assisted breeding efforts. Our history dependent recorder should also function across different non-plant organisms and should be applicable to challenges such as studying differentiation paths involving lowly-expressed genes or evaluating cell to cell variability in gene expression trajectory. In applications beyond tracking gene expression, integrase circuits like the ones described here could be modified for engineering applications. If the reporters were replaced with developmental genes, and integrase expression was tied to chemical signals, specific stress responses, or specific tissues—interactive synthetic genetic programming of development should be entirely feasible for applications such as programming climate-resilient root architectures. The relative ease and accessibility of our method should enable rapid prototyping and optimization for use in elucidating and reprogramming cell state.

## Material and methods

### Construction of plasmids

Our cloning strategy was based on Golden Gate assembly using appropriate spacer and BsaI-HFv2 (NEB) and BbsI-HF (NEB) as the restriction enzymes (Fig. S8). Details on all primers and constructs can be found in Supplementary Data 2 and 3 respectively. Candidate promoter sequences (SPCH: AT5G53210, MUTE: AT3G06120, AHP6: AT1G80100, GATA23: AT5G26930) were amplified from Col-0 genomic DNA (primer list available in Supplementary Data 2) to add specific Golden Gate spacers. After PCR purification, each level0 promoter sequence was cloned using a Zero Blunt PCR Cloning Kit (ThermoFisher Scientific). The PhiC31 integrase sequence was a gift from the Orzeaz lab. The Bxb1 sequence was a gift from the Bonnet lab. Integrases were amplified using primers with golden gate compatible spacers to generate level 0 integrase parts. Constitutive plant promoters and terminators were purchased from Addgene as part of the MoClo Toolbox for Plants^58^. A mutated version of the pPP2AA3 promoter without BsaI sites was ordered from Twist Bioscience. Other level0 fragments were ordered from IDT as Gblocks: the fluorescent proteins NeonGreen-NLS and mtagBFP2-ER and combinations of integrase sites and restriction sites for the construction of various integrase targets. The mCherry sequence was amplified from a constitutively expressed mCherry construct (donated by Jennifer Brophy). For the integrase target level0 sequences, the p35S or pUBQ10 promoter was added by Golden Gate using BbsI sites (Fig. S8).

The construction of level 1 single integrase constructs (constitutive, stomata and lateral root specific) was performed via Golden Gate reaction in the modified pGreenII-Hygr vector containing compatible Golden Gate sites^58^. The construction of integrase targets was performed with the same methods in a modified pGreenII-Kan vector. Construction of level 2 integrase constructs expressing both PhiC31 and Bxb1 for the full recorders was performed by amplifying completed level 1 integrase constructs using primers with golden gate compatible spacers, then performing Golden Gate reactions in the modified pGreenII-Hygr vector containing compatible Golden Gate sites. The addition of an NLS or DST to level 1 and level 2 integrase constructs was performed using PCR-mediated site-directed mutagenesis (NEB Q5 Site-Directed Mutagenesis, Cat #E0554S). Primers for mutagenesis were designed with NEBasechanger (primer list available in Supplementary Data 2). Enzymes for Golden Gate assembly were purchased from New England Biolabs (NEB, Ipswich, MA, USA). PCR were performed using 2X Q5 PCR master mix (NEB) and GoTaq master mix for colony PCR and genotyping (Promega, Madison, WI, USA). Primers were purchased from IDT (Coralville, IA, USA), and DNA fragments from Twist Bioscience (San Francisco, CA, USA) or IDT. Plasmid extraction and DNA purification were performed using Monarch kits (NEB). Sequences were verified with Sanger sequencing by Azenta Life Sciences or Primordium Labs for whole plasmid sequencing. Chemically-competent cultures of the *E. coli* strain DH5αZ1 (laciq, PN25-tetR, SpR, deoR, supE44, Delta(lacZYA-argFV169), Phi80 lacZDeltaM15, hsdR17(rK −, mK +), recA1, endA1, gyrA96, thi-1, relA1) were transformed with plasmid constructs containing kanamycin resistance. Transformed *E. coli* was grown in LB media (LB broth, Miller) with kanamycin (Millipore Sigma, 50 µg/mL).

Sequences are available at the public benchling link (https://benchling.com/cjmaranas/f_/P8Coz4Vw-maranas-et-al-2024-a-history-dependent-integrase-recorder-of-plant-gene-expression-with-single-cell-resolution-/). Constructs would be available on Addgene.

### Plant growth conditions

*Arabidopsis* seedlings were sown in 0.5 X Linsmaier and Skoog nutrient medium (LS) (Caisson Laboratories) and 0.8% w/v Phyto agar (PlantMedia/bioWORLD), stratified at 4 °C for 2 days, and grown in constant light at 22 °C.

### Construction and selection of transgenic Arabidopsis lines

*Agrobacterium tumefaciens* strain GV3101 was transformed by electroporation, and subsequently grown in LB media with rifampin (Millipore Sigma, 50 μg/mL), gentamicin (Millipore Sigma, 50 μg/mL), and kanamycin (Millipore Sigma, 50 μg/mL). The floral dip method^59^ was used to generate integrase target lines in Col-0, and then transformation of the integrase constructs into the target lines were performed via an adjusted dual dipping approach wherein plants are dipped on two occasions approximately a week apart (with the first dipping occurring when the stems are around 3 inches tall) and the agrobacterium were incubated with MMA (10 mM MgCl2, 10 mM MES pH 5.6, 100 µM acetosyringone) for an hour prior to dipping, as acetosyringone is known to improve agrobacterium-mediated transformation efficiency in *Arabidopsis* explants^60^. For T1 selection: 150 mg of T1 seeds (∼2500 seeds) were sterilized using 70% ethanol and 0.05% Triton-X-100 and then washed using 95% ethanol. Seeds were resuspended in 0.1% agarose and spread onto 0.5X LS Phyto selection plates, using 25 μg/mL of kanamycin for target lines or 25μg/mL kanamycin and 25 μg/mL hygromycin for lines with both the integrase construct and the target. The plates were stratified at 4 °C for 48 h then grown for 7-8 days. To select transformants, tall seedlings with long roots and a vibrant green color were picked from the selection plate with sterilized tweezers and transferred to a new 0.5X LS Phyto agar plate for characterization. Selected *Arabidopsis* seed lines will be made available through the Arabidopsis Biological Resource Center at Ohio State University.

### Plant Genotyping

For genotyping of plants, leaves from approximately 20 day old plants were used. For plant DNA extraction, leaf sections approximately 1 cm^2^ in area were frozen on dry ice and ground for 1 minute into a fine powder. 10μm 0.5M NaOH was added to each sample before boiling them for 1 minute and then adding 100μL neutralization solution (1 part 1M Tris-OH/HCl, pH 8.0 and 4 parts 0.1M TE, pH 8.0). Genotyping PCRs were set up using GoTaq master mix (Promega, Madison, WI, USA), making sure to gently stir the DNA extraction with the pipette tip before pipetting into the reaction.

### Imaging of reporter and integrase lines

Initial screening of root and leaf tissue was performed using a Leica Biosystems microscope (model: DMI 3000) with a 10x objective for imaging roots and a 40x objective for imaging leaf tissue. Further characterisation and images in the manuscript were obtained via confocal imaging.

Confocal imaging of the seedling root and leaf tissue was performed using a Nikon A1R HD25 laser scanning confocal microscope with a Plan Apochromat Lambda 20x objective. Three channels were used: 561 laser and 578-623 detector for RFP imaging; 488 laser and 503-545 detector for GFP imaging; 405 laser and 419-476 detector for BFP imaging. For each image, a Z-stack was recorded. All images were processed using ImageJ FIJI.

#### Confocal imaging of lateral root recorder

After characterization of T1 seedlings and subsequent generation of T2 seeds, 10 day old T2 seedlings were mounted on slides with water and with Parafilm edges to prevent the coverslip from pressing on the root. The main root of each seedling was scanned for pre-emergence lateral roots to image. The gain was set at 75, 45, and 40 respectively for the BFP, RFP, and GFP channels. The laser power was set to 3, 10, and 4, respectively.

#### Confocal imaging of stomata single switches and recorder

After characterization of T1 seedlings and subsequent generation of T2 seeds, the first true leaves from 12-16 day old T2 seedlings were cut off and mounted on slides with 50% glycerol for imaging of the abaxial side of the leaf. The edges of the coverslip were painted with clear nail polish to prevent movement of the sample during imaging. The leaves were scanned to locate the flattest areas to take images. The gain was set at 75, 50, and 35 respectively for the BFP, RFP, and GFP channels. The laser power was set to 2, 12, and 3, respectively.

### Characterization of history-dependent recorder in Arabidopsis transgenic lines

T1 seedlings for each line were grown 4–5 days after transformant selection. Each selected seedling was imaged at 10X magnification using an epifluorescence microscope (Leica Biosystems, model: DMI 3000) using the GFP (exposure 400 ms, gain 2), RFP (exposure 500 ms, gain 2), and CFP (exposure 300 ms, gain 2) channels. Selected T1 seedlings were then transferred to soil, and at maturation T2 seeds were collected. For later generations, seedlings were sterilized similarly to T1s, stratified, plated on an LS agar plate, grown for either 10 days (for characterizing roots) or 12-16 days (for characterizing leaves). Target characterization was done using the epifluorescence microscope as for T1. For the target lines, the seedlings with the highest level of mtagBFP2 expression in the root (or in the case of the pre-switched target the highest level of mCherry expression) were selected and transferred to soil to generate T2 seeds. The line with the most consistently bright seedlings was maintained as the target line for each integrase target and used for all later transformations of integrase constructs.

For the constitutive integrase constructs in a target line, around 5-10 T1 seedlings were analyzed per construct and the ones which displayed any level of switching were transplanted to soil for characterization in the T2 generation where around 20 seedlings were characterized per line. Representative root images were taken using the RFP, GFP and CFP channels and merged for final images. Representative leaf images were generated using a Nikon A1R HD25 laser scanning confocal microscope as described above.

### Characterization in the context of lateral root development

For the pAHP6 and pGATA23 single integrase switches, at least 15 T1 seedlings were analyzed per construct. Each seedling was categorized into one of three classes based on specificity of switching: ‘as expected’ (RFP expression in xylem pole, XPP, and lateral root cells); overswitched (RFP expression in additional cells); underswitched (RFP expression in fewer cells). The seedlings in the no switch category represented around 25% of seedlings (Source data 1) and were omitted from analysis as this proportion is consistent with a loss of the integrase construct through gene segregation^21^. Representative images were taken in the epifluorescence microscope using the GFP, RFP, and CFP channels and merged for final images. For the *AHP6* switch, T1 seedlings which showed switching in the xylem poles, xylem pericycle, and lateral root cells were transplanted to soil along with T1 seedlings which showed switching only in the lateral root. For the pGATA23 single integrase switch, only T1 seedlings which showed switching specific to the lateral root were transplanted for future T2 characterization. For each selected T1 line, 15-20 T2 seedlings were characterized, categorizing each as follows: ‘as expected’ (RFP expression in lateral root cells); overswitched (RFP expression in additional cells); underswitched (RFP expression in fewer cells) (Table S1).

For the full lateral root recorder to track both *AHP6* and *GATA23* expression, nine T1 seedlings were analyzed and those whose switch pattern was either ‘as expected’ or some form of underswitched (Fig. S5), were transplanted to soil for future T2 characterization. From each T1 line, 15-20 seedlings were characterized and categorized based on switch pattern (Fig. S6). For Fig. 2, these categories were consolidated into underswitched, ‘as expected’, and overswitched categories (Table S1) as for the single switches, with any seedlings with Bxb1 and/or PhiC31 overswitching fitting into the overswitched category and any seedlings with Bxb1 and/or PhiC31 underswitching fitting into the underswitched category (Fig. S6). Representative images for the full lateral root recorder were taken with the Nikon A1R HD25 laser scanning confocal microscope as described previously (source data 2).

### Characterization in the context of stomata development

For the single pSPCH and pMUTE T1 single integrase switches, leaves from at least 10 T1 seedlings were analyzed per construct. Due to the chlorophyll autofluorescence causing difficulties imaging mCherry in the epifluorescence microscope, this characterization was done using the confocal microscope. For the pSPCH single integrase switch, T1 seedlings with switching in only stomata and some epidermal cells which border stomata were transplanted for T2 characterization. T1 seedlings with switching limited to only stomata were also transplanted. For the pMUTE single integrase switch, seedlings with switching only in the stomata were transplanted for T2 characterization. For the full stomata recorder for tracking *SPCH* and *MUTE* expression, nine T1 seedlings (Table S3) were characterized and any with GFP expression specific to stomata were transplanted for T2 characterization.

For the pSPCH and pMUTE single integrase switch T1 lines, leaves from 15-20 seedlings were characterized as described for T1. The pSPCH single integrase switch seedlings were sorted into one of three categories: ‘as expected’ (RFP expression in guard cells and surrounding epidermal cells); overswitched (RFP expression in additional cells); underswitched (RFP expression in fewer cells). For the pMUTE single integrase switch, the categories are as follows: ‘as expected’ (RFP expression in only guard cells); overswitched (RFP expression in additional cells); underswitched (RFP expression in fewer cells). For the full recorder the categories are as seen in Fig. S7. For Fig. 3 these categories were consolidated into the same underswitched, ‘as expected’, and overswitched categories as for the single switches, with seedlings with either or both PhiC31 and Bxb1 overswitching fitting into the overswitched category and seedlings with either PhiC31 or Bxb1 underswitching fitting into the underswitched category. Representative images for the single switches and the full recorder were taken with the Nikon A1R HD25 laser scanning confocal microscope as described previously (source data 2).

For quantifying the relationship between recorder output and stomata type, leaves from seedlings were stained with 2mg/ml Calcofluor White for 30 minutes (Millipore Sigma Cat #18909) to better visualize the cell boundaries. To screen random stomata all across the leaf, we started from the top left of each leaf and shifted the viewing frame of the microscope across the leaf incrementally and then down at least a full frame before moving back across the leaf. In each frame, the centermost stomata was chosen to be categorized. This categorization involved determining the type of stomata based on the pattern of surrounding cells and then noting the recorder output (GFP expressing or no fluorescence). In the case that the type of stomata was not able to be determined, the frame was skipped. This process continued for each seedling until at least 15 A and 15 NA stomata were counted.

### Analysis

For each single switch construct, the percentage of seedlings in each of the three categories were plotted in a bar plot with the number of seedlings tested mentioned at the top of the bar. For the full recorders the percentage of seedlings in each of the expanded seven categories (Fig. S5, Fig. S6) was plotted but consolidated into the same three categories for display in Figures 2 and 3. Data available in source data file 1.

For quantifying the relationship between stomata type and recorder output, the percentage of State 2 and State α stomata of each type was plotted. For comparisons between A and NA stomata, a student’s t test was performed to evaluate statistical significance.

Python data analysis script which includes statistical tests and plotting functions was run in version 3.9.1 and with the following package dependencies: pandas (version 1.5.3), scipy.stats (version 1.10.0), matplotlib.pyplot (version 3.6.3), matplotlib.colors (version 3.6.3), scikit_posthocs (version 0.21), seaborn (v0.12.0), and numpy (version 1.24.2). All the codes are available in github: https://github.com/cmaranas/Maranas-et-al.-2024-A-history-dependent-integrase-recorder-of-gene-expression.git

All images taken during seedling characterization were opened and processed using the ImageJ FIJI program (version 1.53c). Each .tif image file contained the images of a seedling’s GFP, RFP, and CFP channels. tif files were processed with adjustments to the color lookup table, brightness, and contrast of each channel (GFP: Green, Min 200, Max 2500) (RFP: Red, Min: 300, Max: 3000) (CFP: Blue, Min: 100, Max: 3000).

## Acknowledgement

We thank Janet Solano Sanchez, Ben Downing, and Dr. Alexander Leydon, as well as other members of the Nemhauser, Imaizumi, Di Stilio, Steinbrenner, and Patron groups, for feedback and discussion. We thank Eric Yang for developing and gifting the pPP2AA3 promoter, the Orzáez lab for the PhiC31 sequence, the Brophy lab for the mCherry sequence, and members of the Bonnet lab for sending us the Bxb1 integrase plasmid. We thank Jonah C. Chu for the construction of early versions of the targets and integrase constructs. We thank Keiko Torii for discussions and advice about stomatal development. This work was supported by grants from the National Institutes of Health (grant no. GM107084), the National Science Foundation (grant no. IOS-1546873), and the Howard Hughes Medical Institute Faculty Scholars Program. In addition, support to S.G. was provided by UK Research and Innovation (UKRI) Biotechnology and Biological Sciences Research Council (BBSRC) via the Earlham Institute Core Capability Grant BB/CCG2220/1.

## Author contributions

C.M., S.G. and J.L.N. designed the project. C.M., W.G., and S.G. designed the constructs. C.M., W.G., and S.S. generated the constructs. C.M., S.S., and S.V. performed and analyzed the constitutive integrase switching experiments, C.M. performed and analyzed the lateral root and stomata differentiation experiments, C.M., S.G., and J.L.N. wrote the manuscript.

---

**Figure S1:**
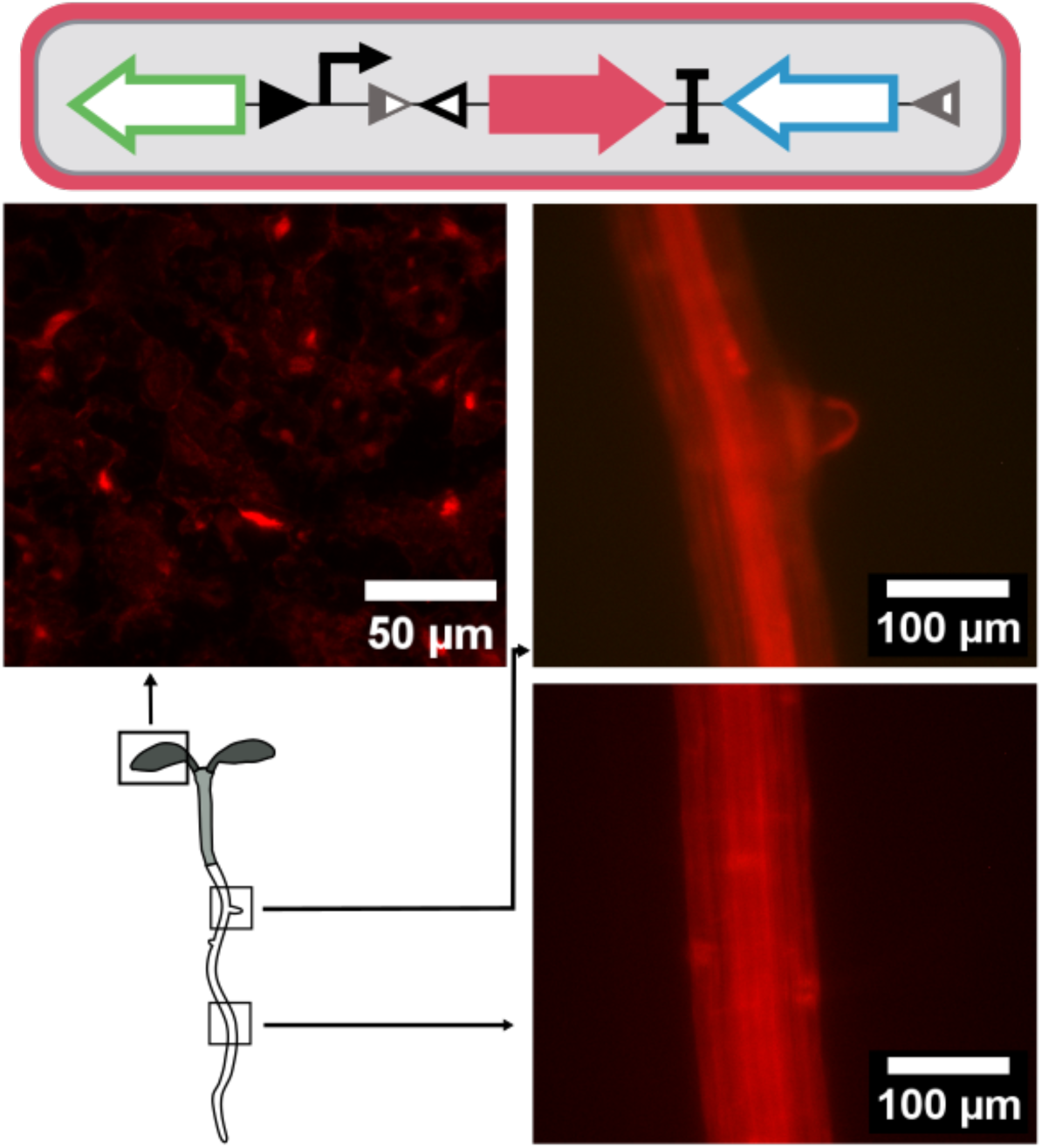
Preswitched PhiC31 then Bxb1 integrase target circuit shows strong fluorescent expression in the leaf and root. The preswitched target was designed to test the State 1 to State 2 switch in the PhiC31 then Bxb1 target such that the target construct is initially in State 1, expressing RFP as per the shown target construct schematic (top). Representative leaf and root images of a T2 seedling transformed with the preswitched target (bottom).

**Figure S2:**
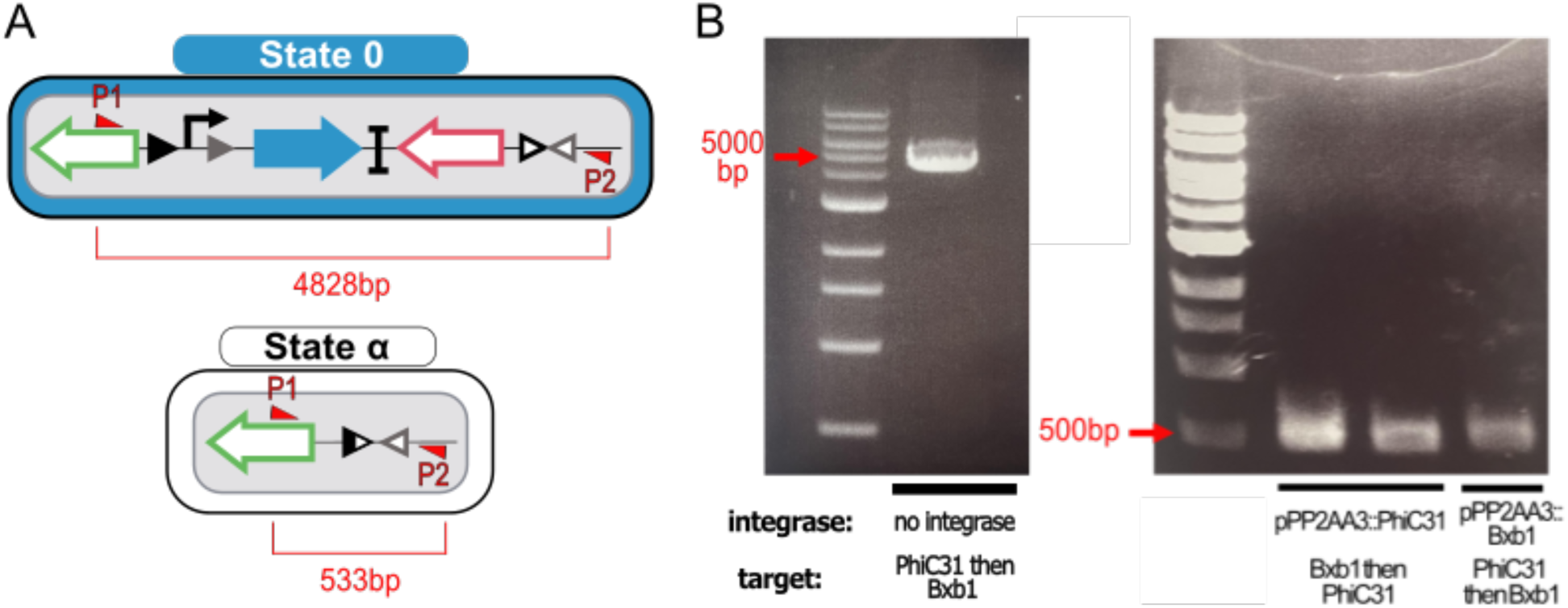
State α genotyping. **A** Schematics of the primer binding on the full length target (shown as State 0, but will be the same size for State 1 or 2) and of the State α target. The PCR of the full length target would result in a band length of 4828 bp whereas pcr of the excised target would result in a band size of 533 bp. **B** Genotyping results. (left) Control PCR for the PhiC31 then Bxb1 full length target in State 0. (right) Genotyping results for pPP2AA3::PhiC31 in the Bxb1 then PhiC31 target (Fig. S3) (left two lanes, corresponding to two different T1 seedlings) and pPP2AA3::Bxb1 in the PhiC31 then Bxb1 target (right lane, one T1 seedling).

**Figure S3:**
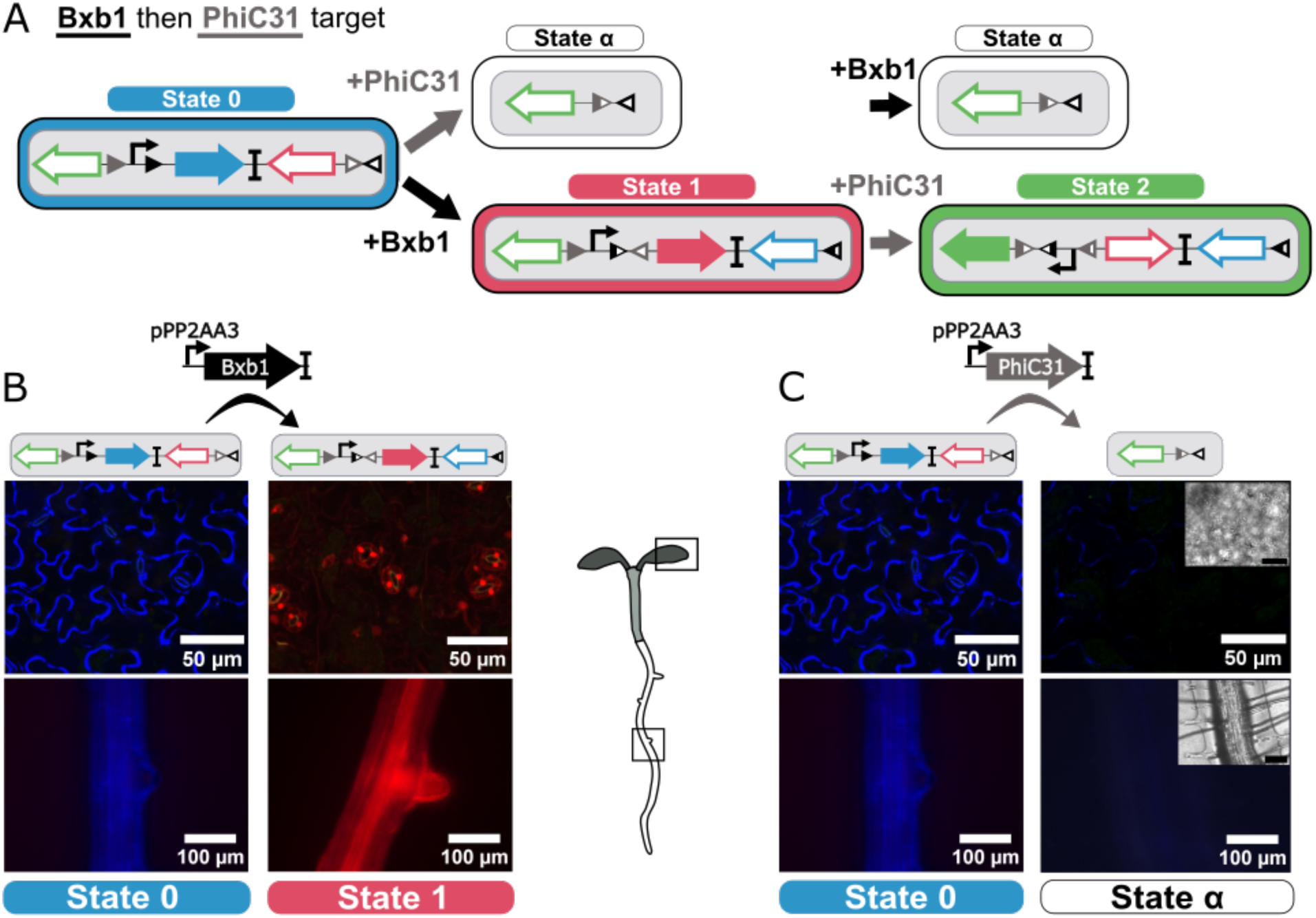
The reverse order history dependent integrase circuit switches between fluorescent states. **A** Schematic of integrase target circuit. The design of the target is the same as in Fig. 1, but Bxb1 mediates the switch from State 0 to 1 and PhiC31 mediates the switch from State 1 to 2. Addition of the PhiC31 integrase to the State 0 target configuration results in a DNA excision and a loss of fluorescence (State α). **B** The Bxb1 integrase mediates switch from State 0 to State 1. (left) The initial target in State 0 shows strong BFP mtagBFP2 expression in the leaf and the root. (middle) Addition of constitutive Bxb1 expression to the target mediates a switch to State 1 and strong RFPmCherry fluorescence in leaf and root tissue. **C** Addition of PhiC31 prior to Bxb1 mediates an “out of order” switch to State α and loss of fluorescence. (left) The initial target in State 0 shows strong BFPmtagBFP2 expression in the leaf and the root. (right) Addition of constitutively expressed PhiC31 to the target in State 0 results in a switch to State α and a loss in fluorescence in the leaf and root. Black scale bars in the brightfield images correspond to 100 μm (root) and 50 μm (leaf).

**Figure S4:**
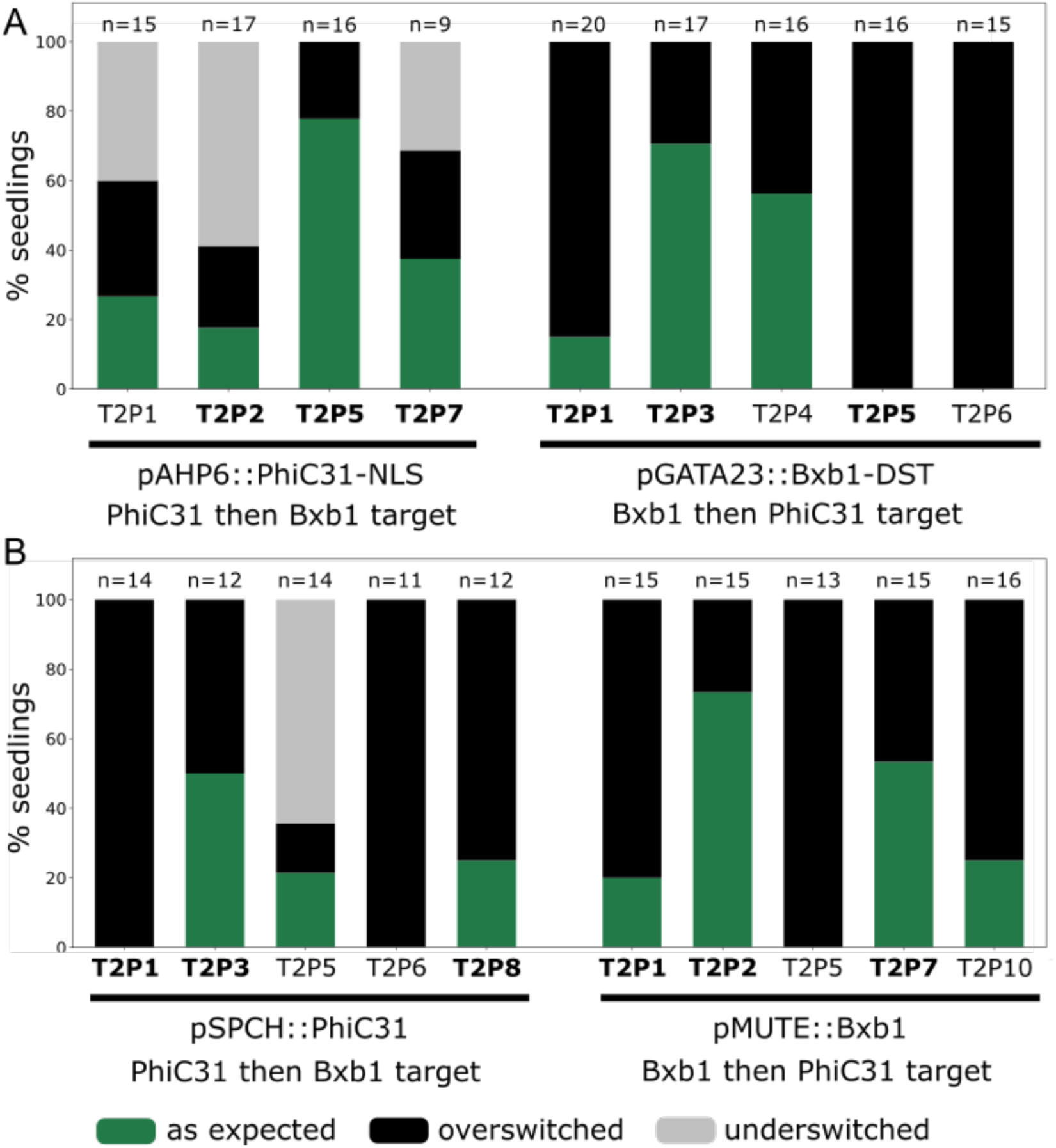
Single integrase switch characterization for lateral root and stomata related genes. The bar graph corresponds to the percentage of seedling per T1 line in each category: ‘as expected’ (in green), overswitched (in black), and underswitched (in light grey). **A** Single integrase switch performance for lateral root genes. Seedlings were screened from T1 lines which came from T1 plants with an ‘as expected’ or underswitched switch pattern. The PhiC31 then Bxb1 target was transformed with the pAHP6::PhiC31-NLS integrase construct (left). The reverse order Bxb1 then PhiC31 target (Fig. S3) was transformed with the pGATA23::Bxb1-DST integrase construct (right). The number of screened seedlings for each T1 line are indicated above the bars. The bolded T1 lines correspond to the lines shown in Figures 2a and 2b. **B** Single integrase switch performance for stomata genes. Seedlings were screened from T1 lines which came from T1 plants with an ‘as expected’ or underswitched switch pattern. The PhiC31 then Bxb1 target was transformed with the pSPCH::PhiC31 integrase construct (left). The Bxb1 then PhiC31 target (Fig. S3) was transformed with the pGATA23::Bxb1 integrase construct (right). The number of screened seedlings for each T1 line are indicated above the bars. The bolded T1 lines correspond to the lines shown in Figures 3a and 3b. Source data is provided as a Source Data file 1.

**Figure S5:**
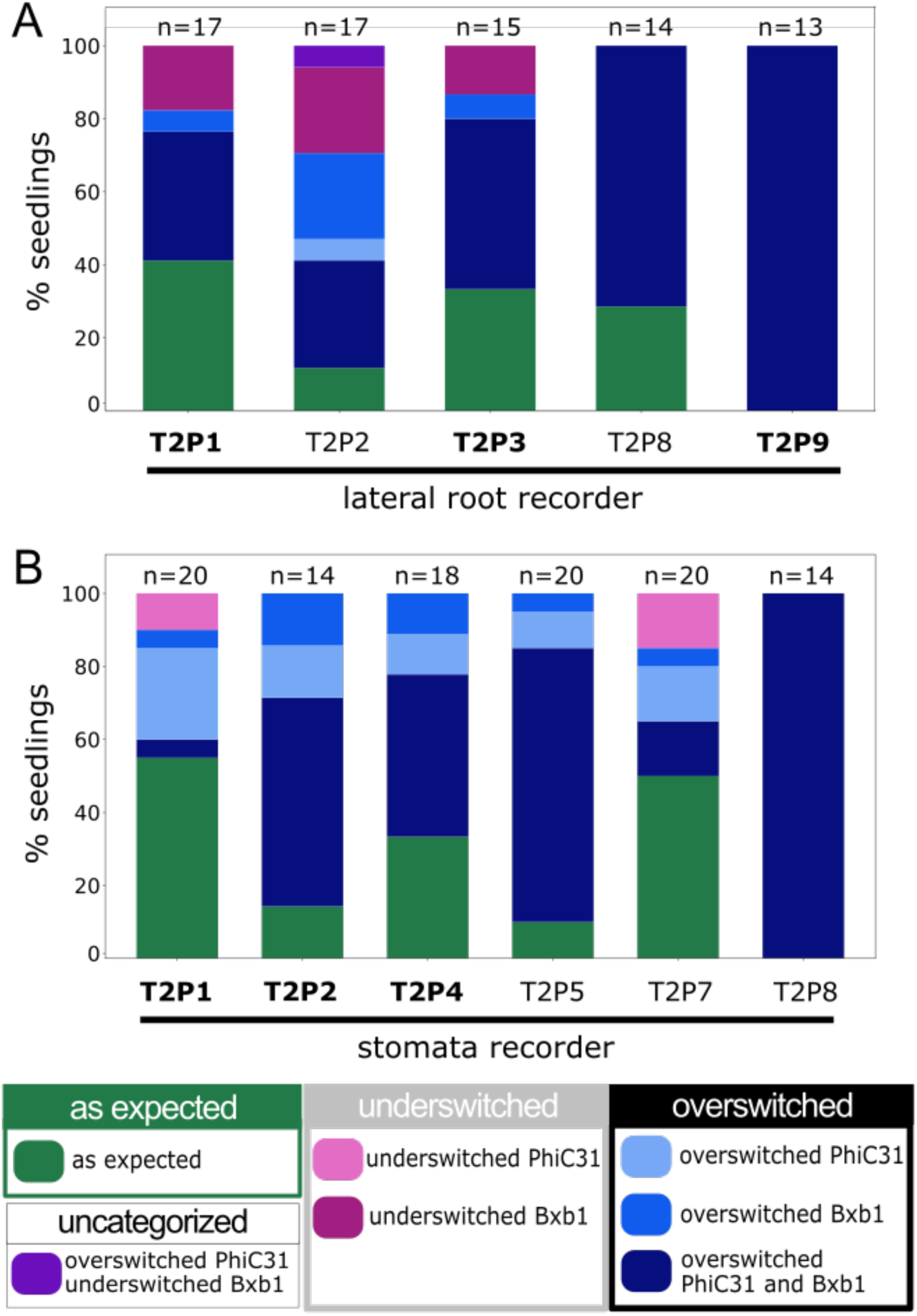
Detailed characterization of recorder performance. The bar graph corresponds to the percentage of seedling per T1 line in each category. **A** Lateral root recorder performance. Seedlings were screened from 5 T2 seedlings from T1 lines which came from T1 plants with an ‘as expected’ or underswitched recorder output. The PhiC31 then Bxb1 target was transformed with the pAHP6::PhiC31-NLS-pGATA23::Bxb1 dual integrase construct. The number of seedlings screened for each T1 line is indicated above the bars. Bolded T1 lines correspond to the lines shown in Fig. 2d. The detailed switch categories correspond to the representations in Fig. S6. **B** Stomata recorder performance. Seedlings were screened from 6 T1 lines which came from T1 plants with an ‘as expected’ or underswitched recorder output. The PhiC31 then Bxb1 target was transformed with the pSPCH::PhiC31-pMUTE::Bxb1 dual integrase construct. The number of seedlings screened for each T1 line is indicated above the bars. Bolded T1 lines correspond to the lines shown in Fig 3d. The detailed switch categories correspond to the representations in Fig. S7. Source data is provided as a Source Data file 1.

**Figure S6:**
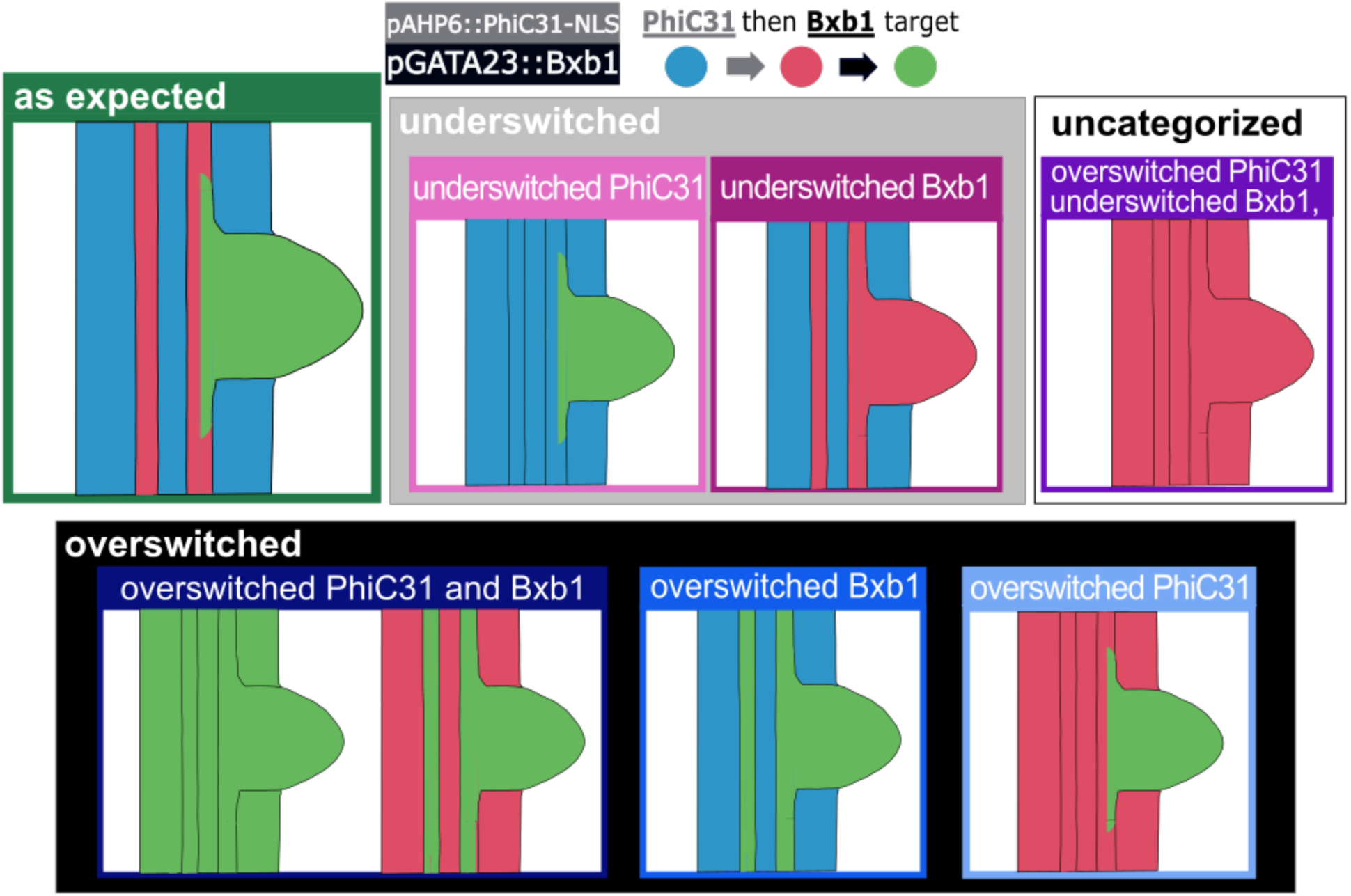
Schematics for observed lateral root recorder output. The diagrams represent the lateral root recorder outputs observed for the PhiC31 then Bxb1 target transformed with the pAHP6::PhiC31-NLS-pGATA23::Bxb1 integrase construct. The box colors correspond to the key in Fig. S5 and the background boxes correspond to the categories in Figure 2.

**Figure S7:**
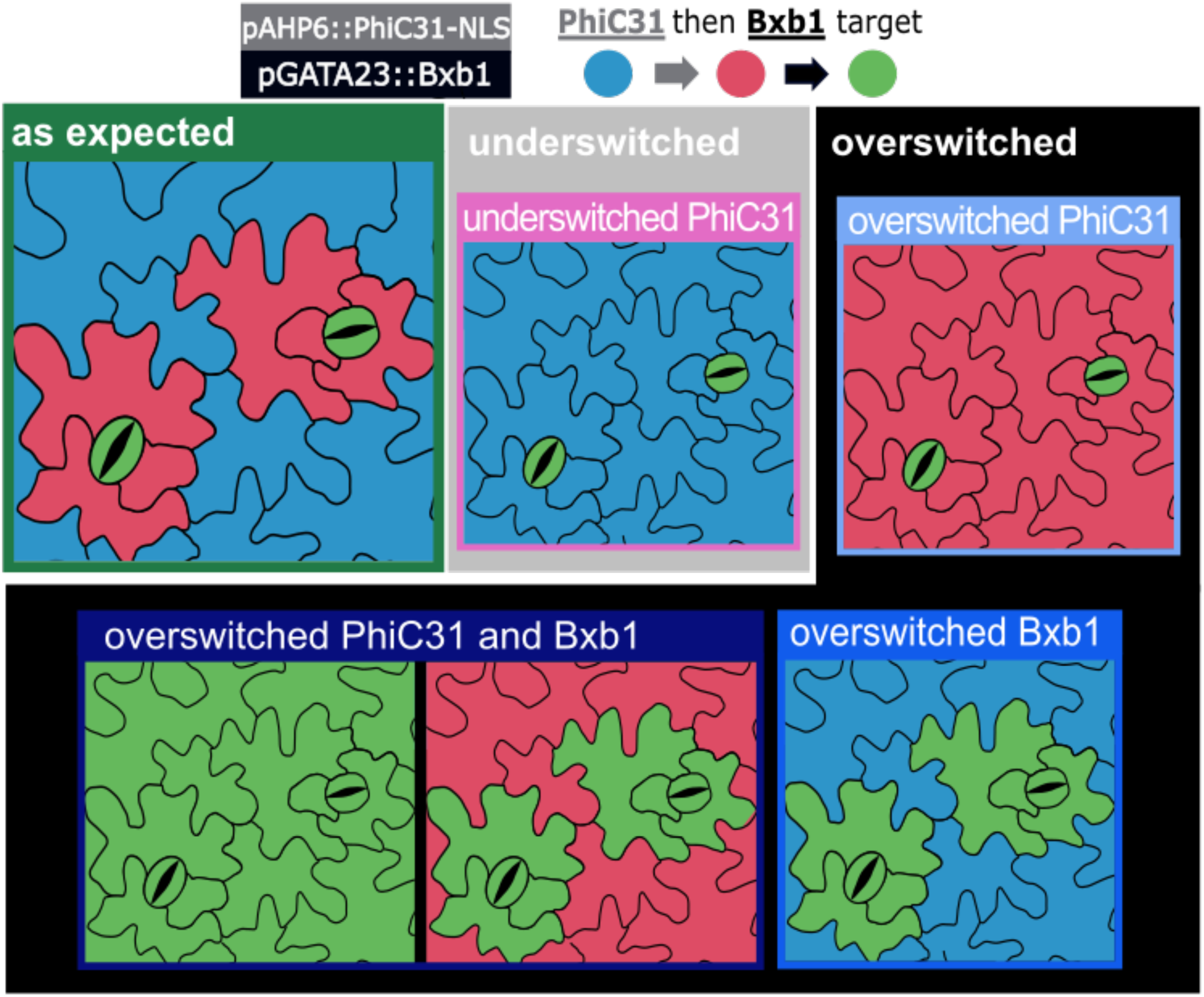
Schematics for observed stomata recorder output. The diagrams represent the lateral root recorder outputs observed for the PhiC31 then Bxb1 target transformed with the pSPCH::PhiC31-pMUTE::Bxb1 integrase construct. The box colors correspond to the key in Fig. S5 and the background boxes correspond to the categories in Figure 3.

**Figure S8:**
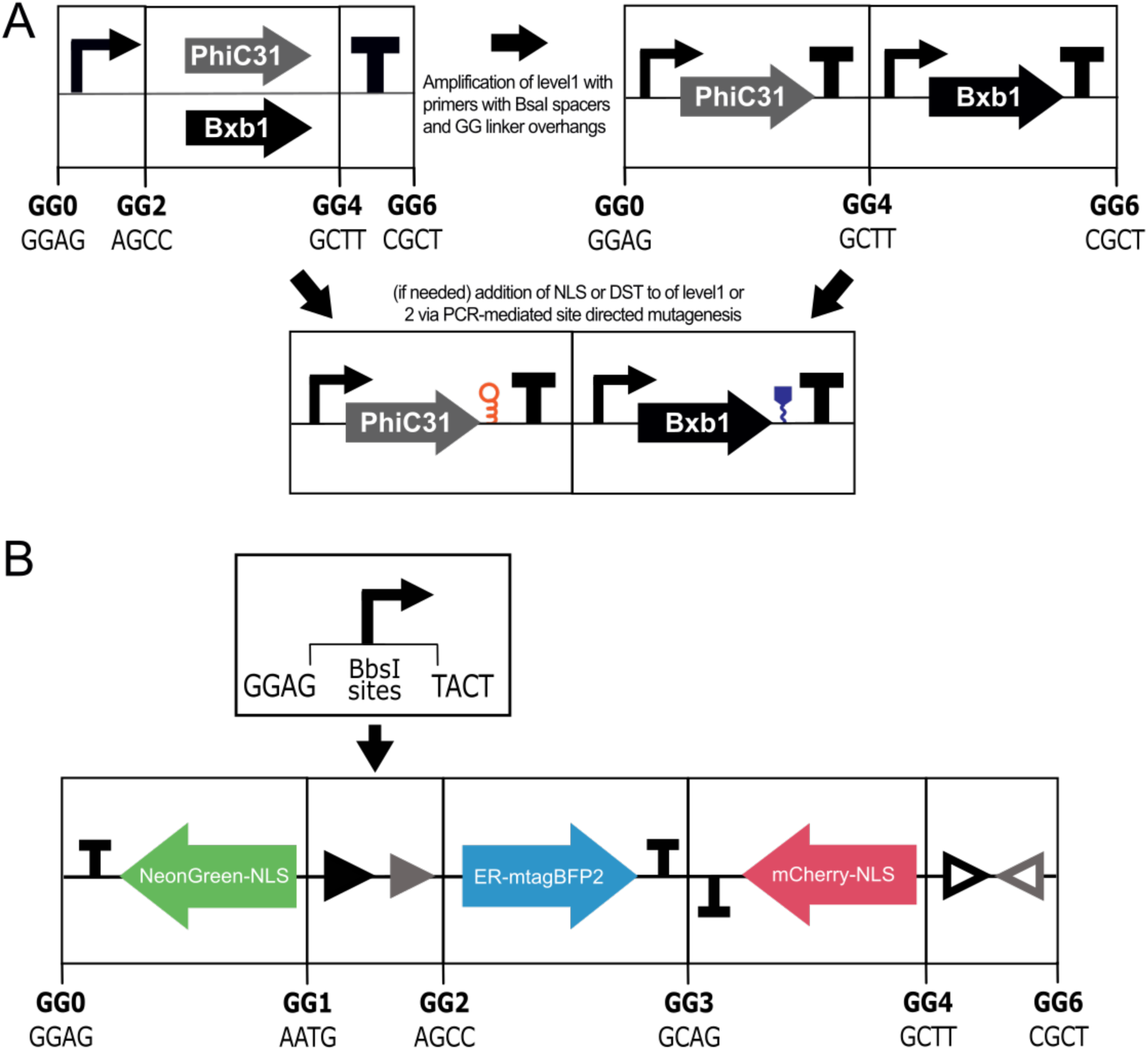
Cloning strategy. **A** Cloning strategy for the integrase level 1 and level 2 constructs. The level 1 is built using golden gate cloning with BsaI restriction sites. The level 1 integrase constructs are then amplified using primers with overhangs including a BsaI restriction site and the appropriate GG sticky end. Then the level 2 is assembled using golden gate assembly with BsaI. If desired, tuning parts (NLS, DST) are added to the level 1 or level 2 via PCR-mediated site directed mutagenesis. **B** Cloning strategy for the target construct. The part with the integrase sites and fluorescent proteins is constructed by golden gate assembly with the BsaI enzyme. The promoter is added to the synthetic fragment via golden gate cloning using BbsI restriction sites to form the completed target.

**Table S1:** Overview of integrase lines and characterization categories.

|  | Integrase construct name | PhiC31 construct | Bxb1 construct | Target construct | As expected | Underswitched | Overswitched |
| --- | --- | --- | --- | --- | --- | --- | --- |
| pAHP6 single switch | L1P137 | pAHP6::PhiC31-NLS:tUBQ1 | n/a | PhiC31 then Bxb1 target (Fig. 1) | RFP in xylem and lateral roots BFP otherwise | RFP in less cells than xylem and LR | RFP in more cells than xylem and LR |
| pGATA23 single switch | L1B42 | n/a | pGATA23::Bxb1-DST:tUBQ10 | Bxb1 then PhiC31 target (Fig. S3) | RFP in LR BFP otherwise | RFP in less cells than LR | RFP in more cells than LR |
| Lateral root history-dependent tracker | L1PB10 | pAHP6::PhiC31-NLS:tUBQ1 | pGATA23::Bxb1-DST:tUBQ10 | PhiC31 then Bxb1 target (Fig. 1) | GFP in LR RFP in xylem BFP otherwise | GFP in less cells than LR and/or RFP in less cells than xylem | GFP in more cells than LR and/or RFP in more cells than xylem and LR |
| pSPCH single switch | L1P139 | pSPCH::PhiC31:tUBQ1 | n/a | PhiC31 then Bxb1 target (Fig. 1) | RFP in guard cells and surrounding epidermal cells BFP otherwise | RFP in less cells than guard cells and surrounding epidermal cells | RFP in more cells than guard cells and surrounding epidermal cells |
| pMUTE single switch | L1B39 | n/a | pMUTE::Bxb1:tUBQ10 | Bxb1 then PhiC31 target (Fig. S3) | RFP in guard cells BFP otherwise | RFP in less cells than guard cells | RFP in more cells than guard cells |
| Stomatal history-dependent tracker | L1PB1 | pSPCH::PhiC31:tUBQ1 | pMUTE::Bxb1:tUBQ10 | PhiC31 then Bxb1 target (Fig. 1) | GFP in guard cells RFP in surrounding epidermal cells BFP otherwise | GFP in less cells than guard cells and/or RFP in less cells than surrounding epidermal cells | GFP in more cells than guard cells and/or RFP in more cells than surrounding epidermal cells |

**Table S2:**
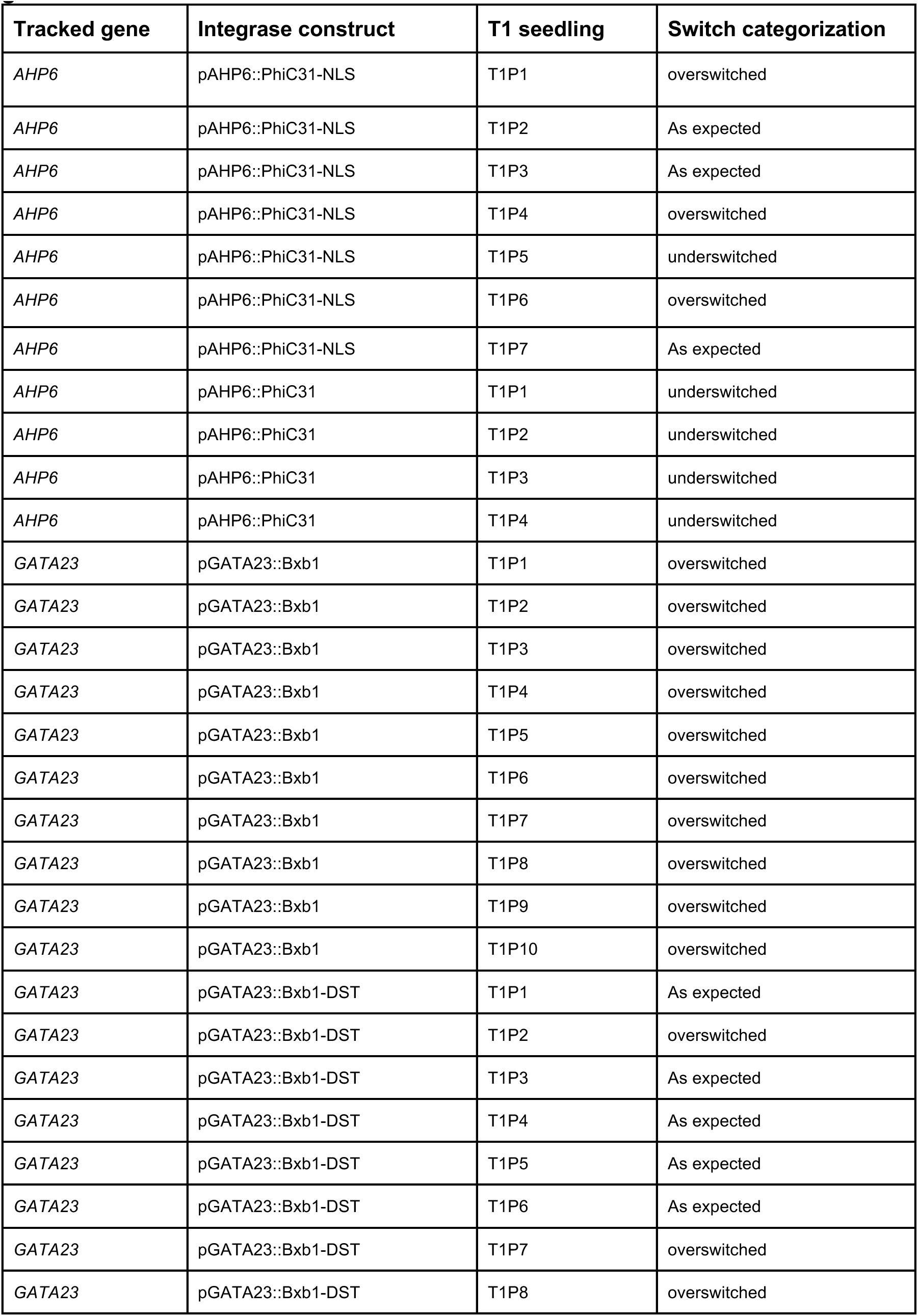
characterization of single integrase switches for lateral root genes in the T1 generation.

| Tracked gene | Integrase construct | T1 seedling | Switch categorization |
| --- | --- | --- | --- |
| *AHP6* | pAHP6::PhiC31-NLS | T1P1 | overswitched |
| *AHP6* | pAHP6::PhiC31-NLS | T1P2 | As expected |
| *AHP6* | pAHP6::PhiC31-NLS | T1P3 | As expected |
| *AHP6* | pAHP6::PhiC31-NLS | T1P4 | overswitched |
| *AHP6* | pAHP6::PhiC31-NLS | T1P5 | underswitched |
| *AHP6* | pAHP6::PhiC31-NLS | T1P6 | overswitched |
| *AHP6* | pAHP6::PhiC31-NLS | T1P7 | As expected |
| *AHP6* | pAHP6::PhiC31 | T1P1 | underswitched |
| *AHP6* | pAHP6::PhiC31 | T1P2 | underswitched |
| *AHP6* | pAHP6::PhiC31 | T1P3 | underswitched |
| *AHP6* | pAHP6::PhiC31 | T1P4 | underswitched |
| *GATA23* | pGATA23::Bxb1 | T1P1 | overswitched |
| *GATA23* | pGATA23::Bxb1 | T1P2 | overswitched |
| *GATA23* | pGATA23::Bxb1 | T1P3 | overswitched |
| *GATA23* | pGATA23::Bxb1 | T1P4 | overswitched |
| *GATA23* | pGATA23::Bxb1 | T1P5 | overswitched |
| *GATA23* | pGATA23::Bxb1 | T1P6 | overswitched |
| *GATA23* | pGATA23::Bxb1 | T1P7 | overswitched |
| *GATA23* | pGATA23::Bxb1 | T1P8 | overswitched |
| *GATA23* | pGATA23::Bxb1 | T1P9 | overswitched |
| *GATA23* | pGATA23::Bxb1 | T1P10 | overswitched |
| *GATA23* | pGATA23::Bxb1-DST | T1P1 | As expected |
| *GATA23* | pGATA23::Bxb1-DST | T1P2 | overswitched |
| *GATA23* | pGATA23::Bxb1-DST | T1P3 | As expected |
| *GATA23* | pGATA23::Bxb1-DST | T1P4 | As expected |
| *GATA23* | pGATA23::Bxb1-DST | T1P5 | As expected |
| *GATA23* | pGATA23::Bxb1-DST | T1P6 | As expected |
| *GATA23* | pGATA23::Bxb1-DST | T1P7 | overswitched |
| *GATA23* | pGATA23::Bxb1-DST | T1P8 | overswitched |

**Table S3:** characterization of the lateral root and stomatal recorder in the T1 generation.

| **Cell lineage** | **Tracked genes** | **PhiC31 construct** | **Bxb1 construct** | **T1 seedling** | **Switch categorization** |
| --- | --- | --- | --- | --- | --- |
| lateral root | *AHP6, GATA23* | pAHP6::PhiC31-NLS | pGATA23::Bxb1 | T1P1 | Underswitched PhiC31 |
| lateral root | *AHP6, GATA23* | pAHP6::PhiC31-NLS | pGATA23::Bxb1 | T1P2 | Underswitched PhiC31 |
| lateral root | *AHP6, GATA23* | pAHP6::PhiC31-NLS | pGATA23::Bxb1 | T1P3 | As expected |
| lateral root | *AHP6, GATA23* | pAHP6::PhiC31-NLS | pGATA23::Bxb1 | T1P4 | Overswitched PhiC31 and Bxb1 |
| lateral root | *AHP6, GATA23* | pAHP6::PhiC31-NLS | pGATA23::Bxb1 | T1P5 | Overswitched PhiC31 and Bxb1 |
| lateral root | *AHP6, GATA23* | pAHP6::PhiC31-NLS | pGATA23::Bxb1 | T1P6 | Overswitched PhiC31 and Bxb1 |
| lateral root | *AHP6, GATA23* | pAHP6::PhiC31-NLS | pGATA23::Bxb1 | T1P7 | Overswitched PhiC31 and Bxb1 |
| lateral root | *AHP6, GATA23* | pAHP6::PhiC31-NLS | pGATA23::Bxb1 | T1P8 | Underswitched PhiC31 |
| lateral root | *AHP6, GATA23* | pAHP6::PhiC31-NLS | pGATA23::Bxb1 | T1P9 | As expected |
| stomata | *SPCH, MUTE* | pSPCH::PhiC31 | pMUTE::Bxb1 | T1P1 | As expected |
| stomata | *SPCH, MUTE* | pSPCH::PhiC31 | pMUTE::Bxb1 | T1P2 | As expected |
| stomata | *SPCH, MUTE* | pSPCH::PhiC31 | pMUTE::Bxb1 | T1P3 | Overswitched Bxb1 |
| stomata | *SPCH, MUTE* | pSPCH::PhiC31 | pMUTE::Bxb1 | T1P4 | As expected |
| stomata | *SPCH, MUTE* | pSPCH::PhiC31 | pMUTE::Bxb1 | T1P5 | As expected |
| stomata | *SPCH, MUTE* | pSPCH::PhiC31 | pMUTE::Bxb1 | T1P6 | Overswitched Bxb1 |
| stomata | *SPCH, MUTE* | pSPCH::PhiC31 | pMUTE::Bxb1 | T1P7 | As expected |
| stomata | *SPCH, MUTE* | pSPCH::PhiC31 | pMUTE::Bxb1 | T1P8 | As expected |
| stomata | *SPCH, MUTE* | pSPCH::PhiC31 | pMUTE::Bxb1 | T1P9 | Overswitched Bxb1 |

**Table S4:**
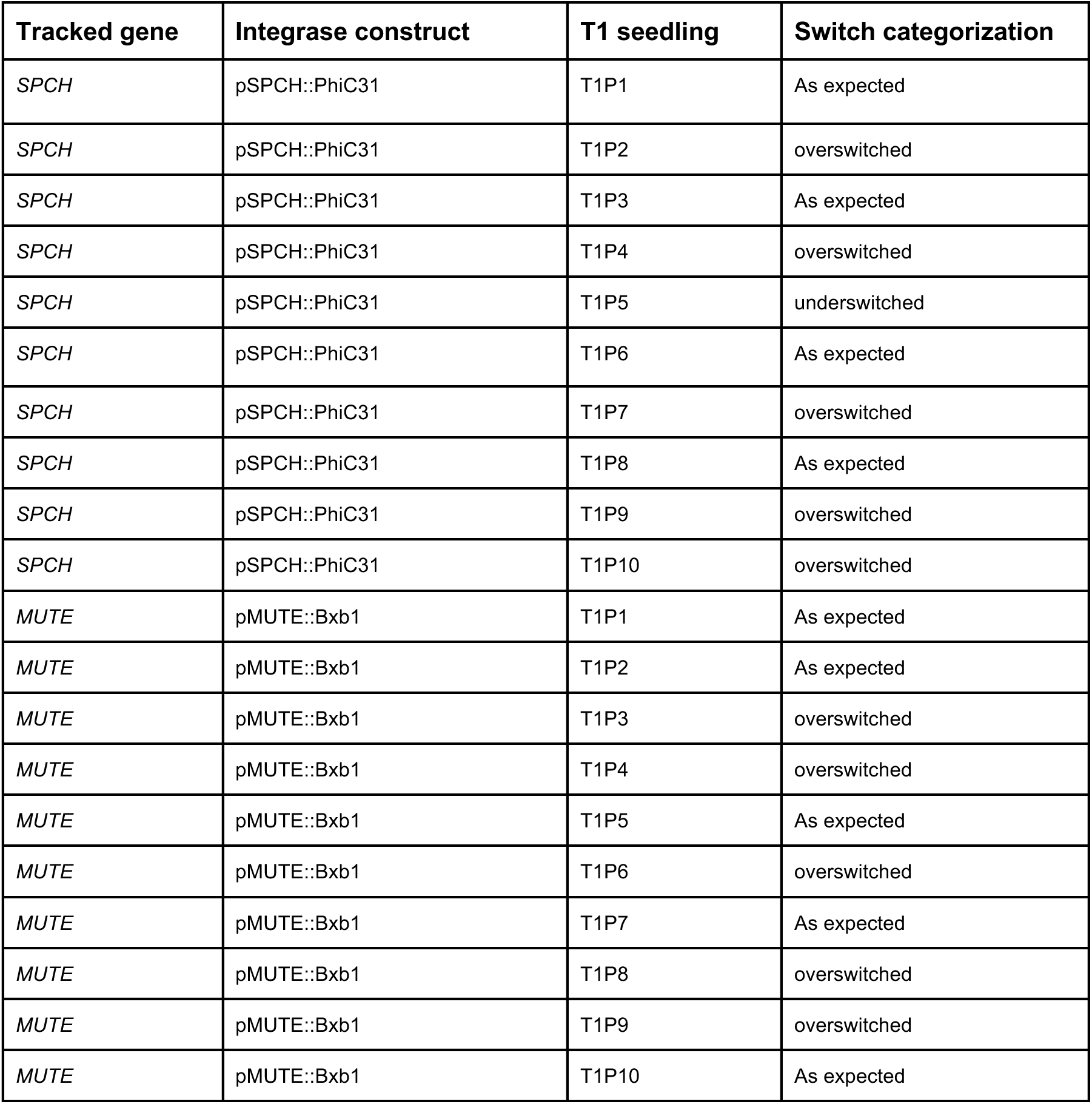
characterization of single integrase switches for stomatal genes in the T1 generation.

| Tracked gene | Integrase construct | T1 seedling | Switch categorization |
| --- | --- | --- | --- |
| *SPCH* | pSPCH::PhiC31 | T1P1 | As expected |
| *SPCH* | pSPCH::PhiC31 | T1P2 | overswitched |
| *SPCH* | pSPCH::PhiC31 | T1P3 | As expected |
| *SPCH* | pSPCH::PhiC31 | T1P4 | overswitched |
| *SPCH* | pSPCH::PhiC31 | T1P5 | underswitched |
| *SPCH* | pSPCH::PhiC31 | T1P6 | As expected |
| *SPCH* | pSPCH::PhiC31 | T1P7 | overswitched |
| *SPCH* | pSPCH::PhiC31 | T1P8 | As expected |
| *SPCH* | pSPCH::PhiC31 | T1P9 | overswitched |
| *SPCH* | pSPCH::PhiC31 | T1P10 | overswitched |
| *MUTE* | pMUTE::Bxb1 | T1P1 | As expected |
| *MUTE* | pMUTE::Bxb1 | T1P2 | As expected |
| *MUTE* | pMUTE::Bxb1 | T1P3 | overswitched |
| *MUTE* | pMUTE::Bxb1 | T1P4 | overswitched |
| *MUTE* | pMUTE::Bxb1 | T1P5 | As expected |
| *MUTE* | pMUTE::Bxb1 | T1P6 | overswitched |
| *MUTE* | pMUTE::Bxb1 | T1P7 | As expected |
| *MUTE* | pMUTE::Bxb1 | T1P8 | overswitched |
| *MUTE* | pMUTE::Bxb1 | T1P9 | overswitched |
| *MUTE* | pMUTE::Bxb1 | T1P10 | As expected |

