## Supplementary material for "A history-dependent integrase recorder of plant gene expression with single cell resolution": Source Data 2

**Source data 2 – Representative microscope images**

Maranas et al.

### pAHP6-NLS single integrase switch (PhiC31 then Bxb1 target transformed with pAHP6::PhiC31-NLS)

**'as expected'**

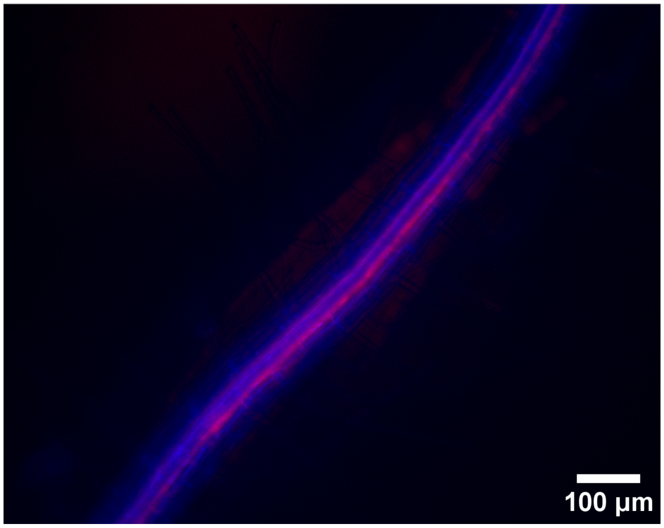


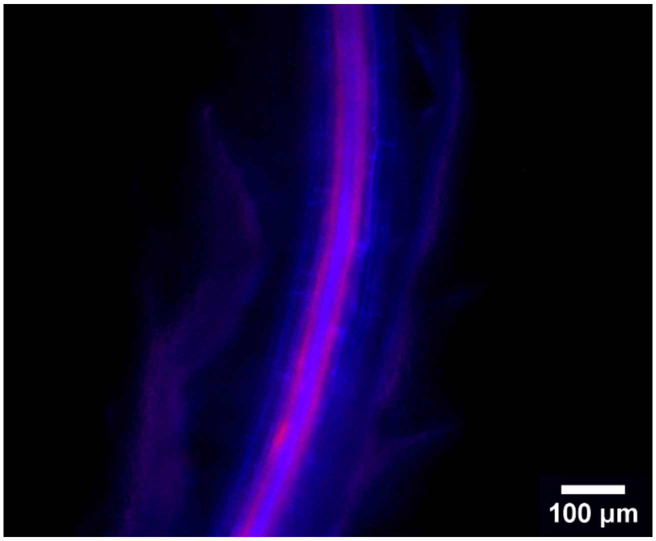


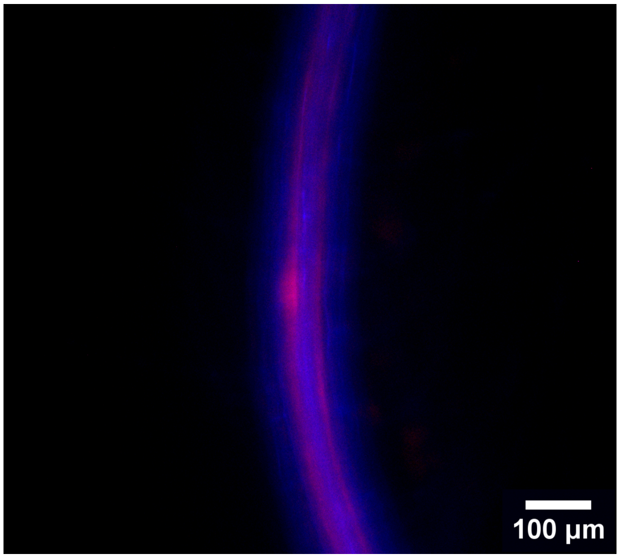

**Underswitched**

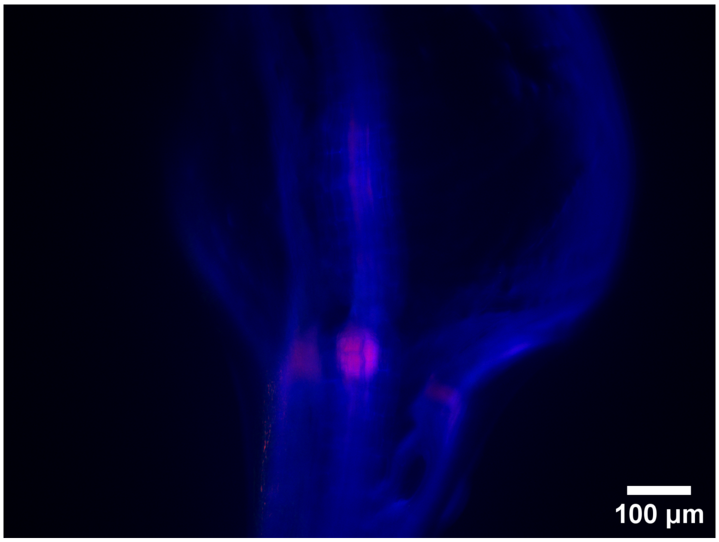


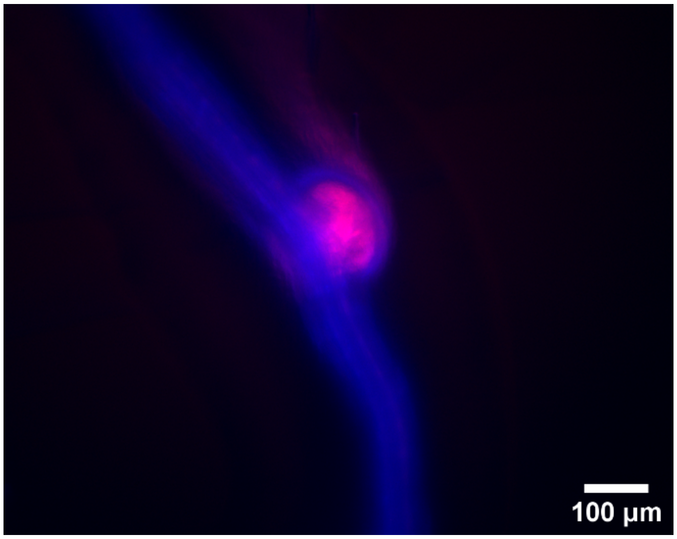


**Overswitched**

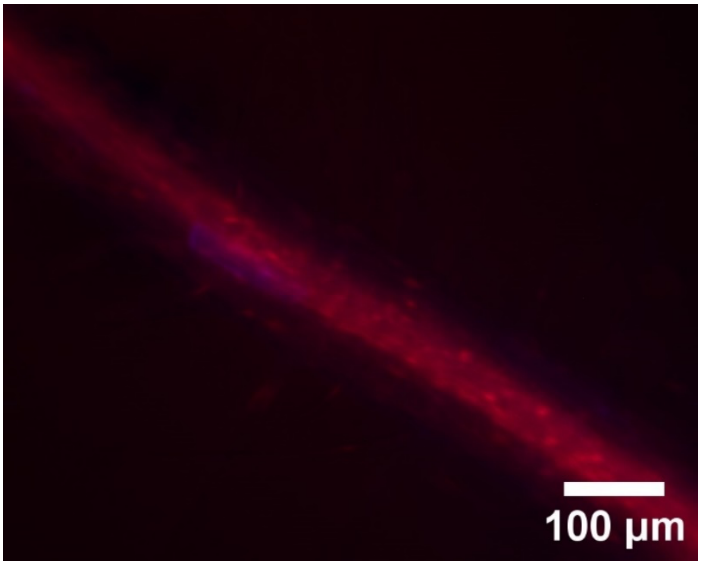


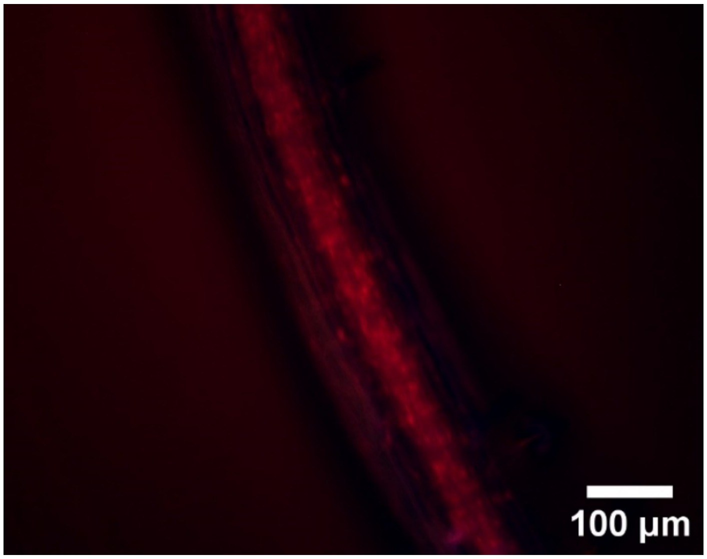

### pGATA23-DST single integrase switch (Bxb1 then PhiC31 target transformed with pGATA23::Bxb1-DST)

**‘as expected’**

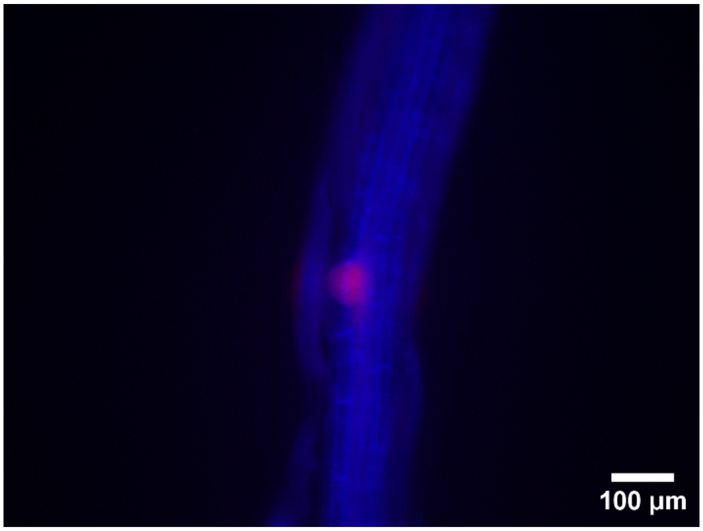


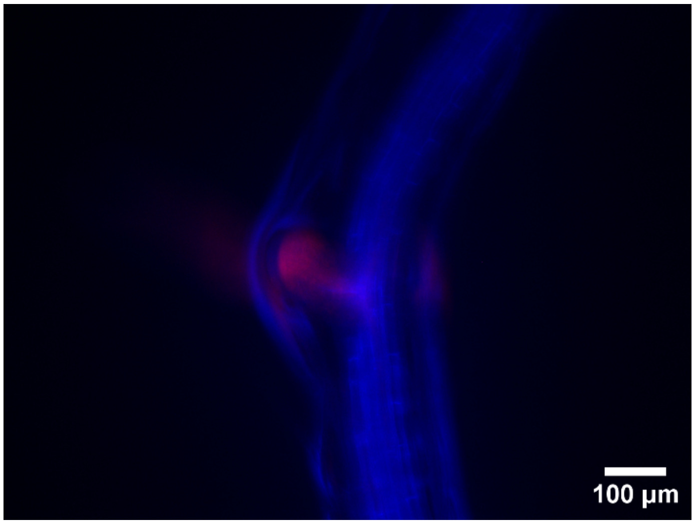


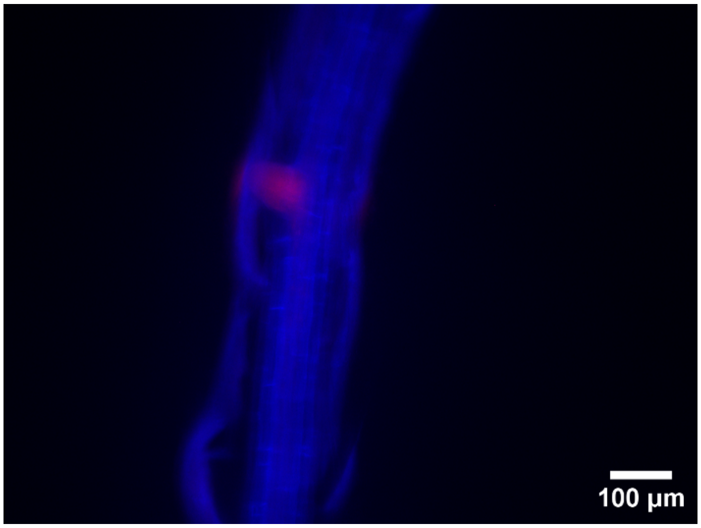


**Overswitched**

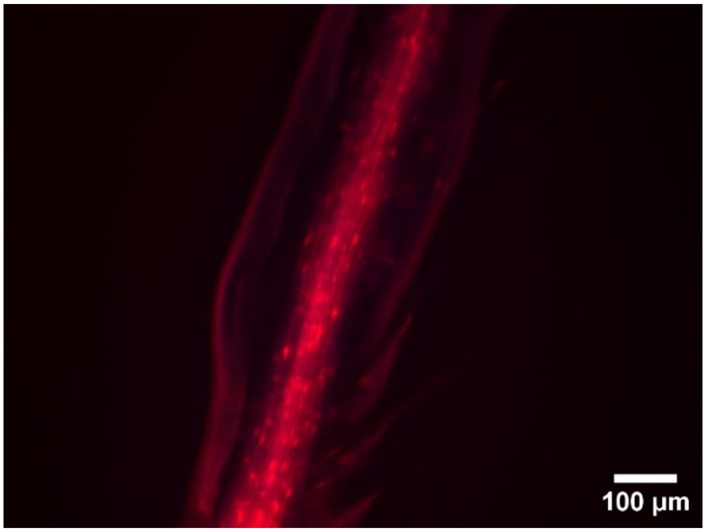


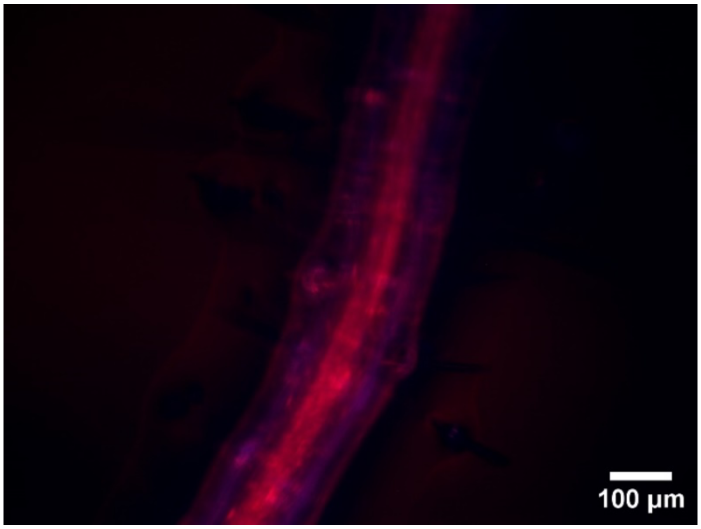

### Full lateral root recorder (pAHP6::PhiC31-NLS, pGATA23::Bxb1 transformed into the PhiC31 then Bxb1 target)

**‘as expected'**

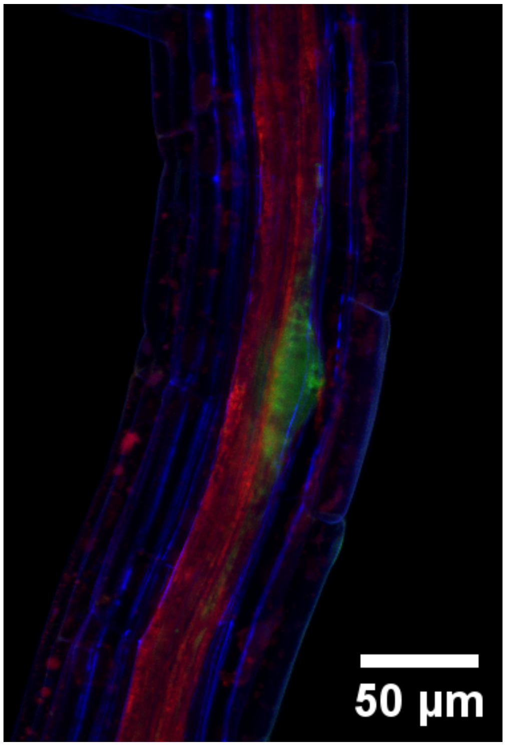


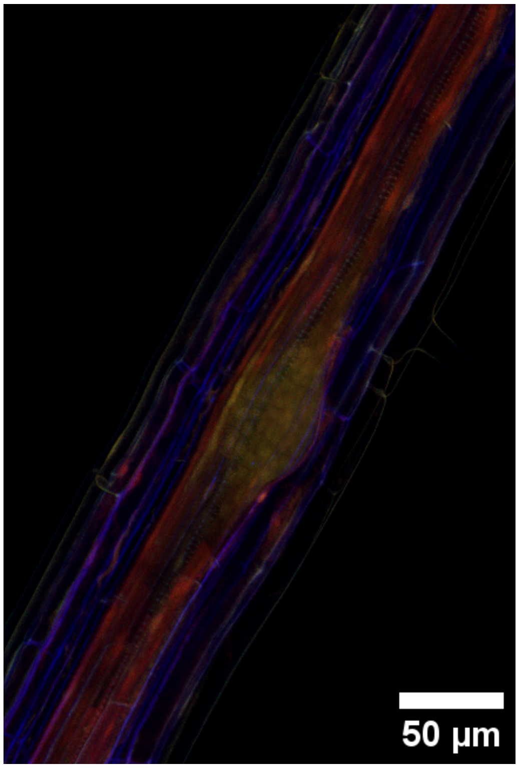


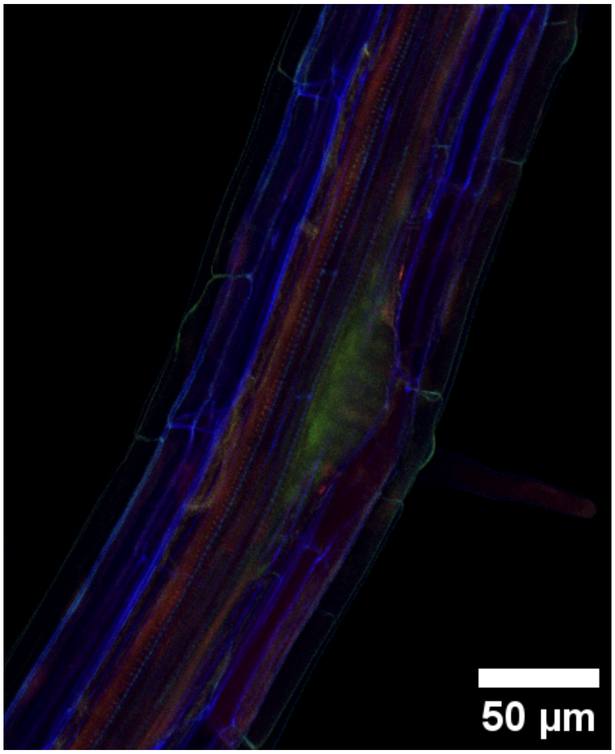

**Bxb1 overswitched**

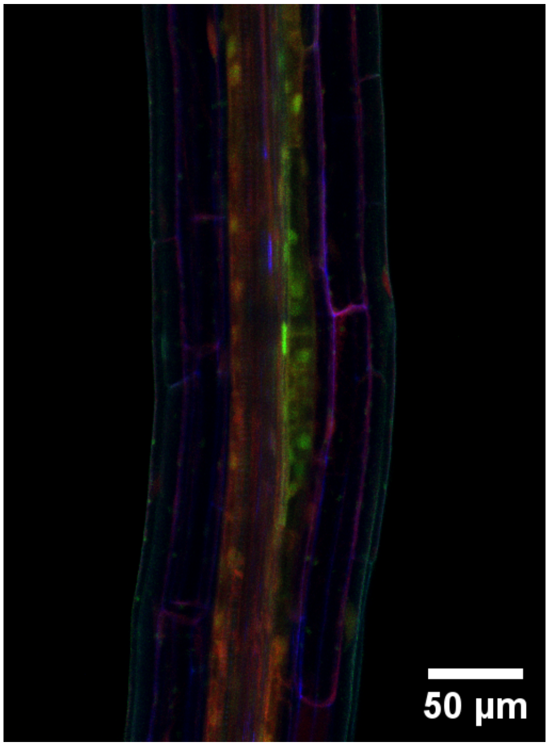


**PhiC31 overswitched**

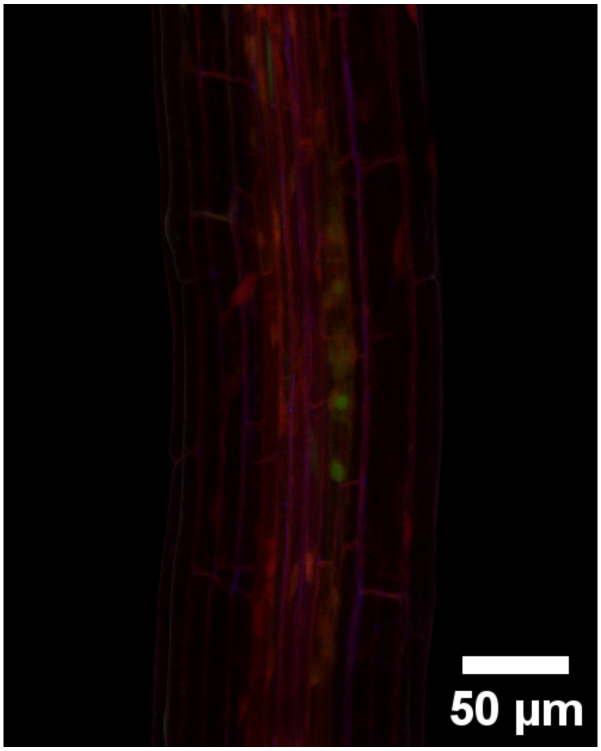


**PhiC31 and Bxb1 overswitched**

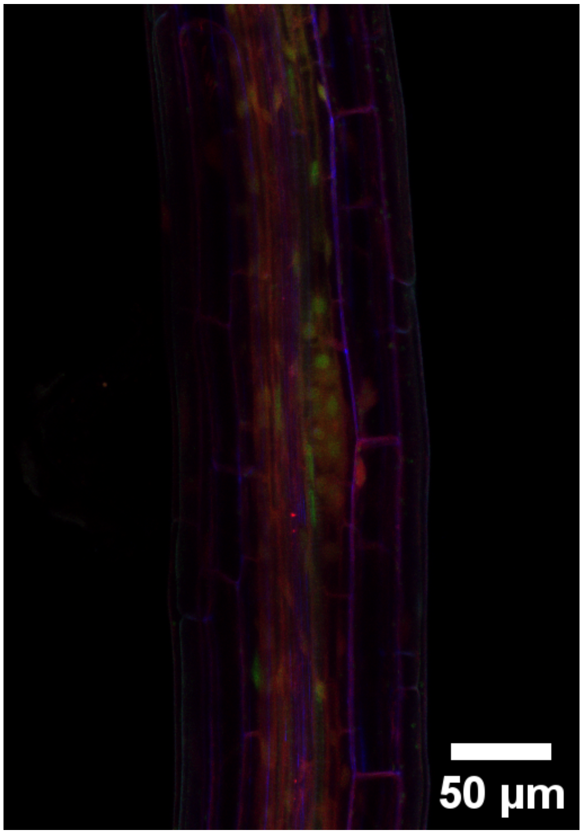


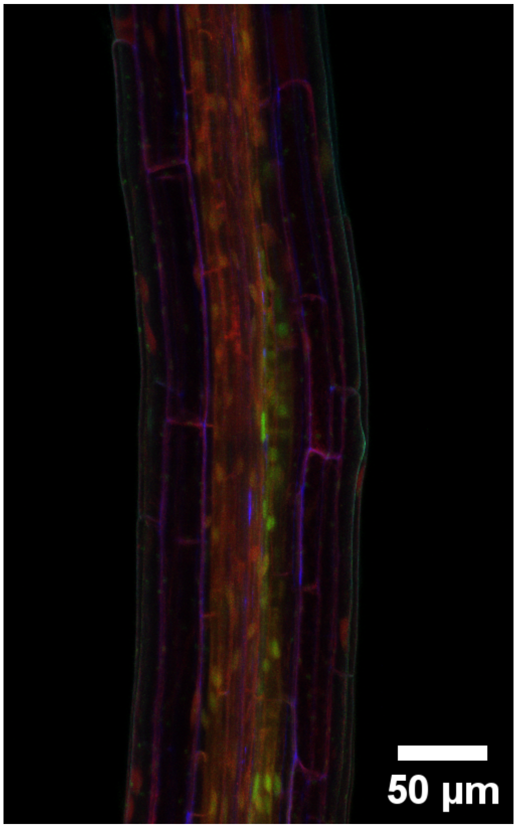

### pSPCH single integrase switch (PhiC31 then Bxb1 target transformed with pSPCH::PhiC31)

**‘as expected’**

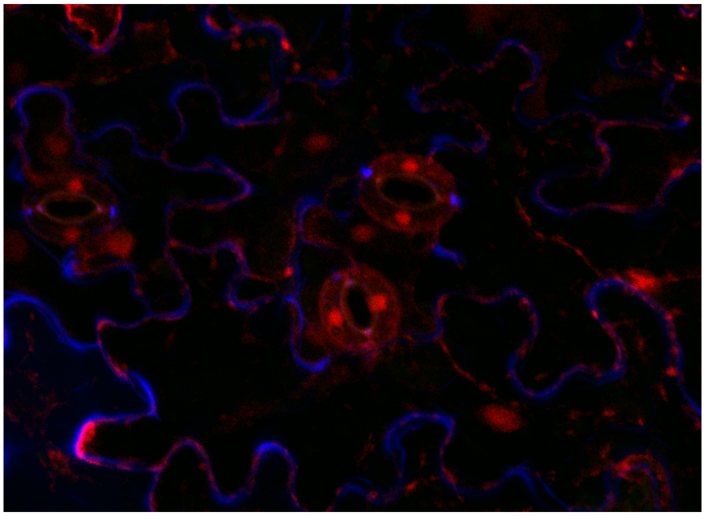

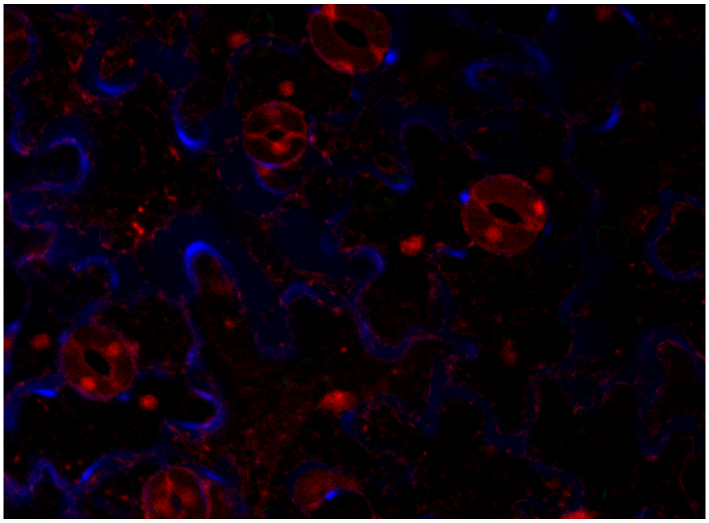

**overswitched**

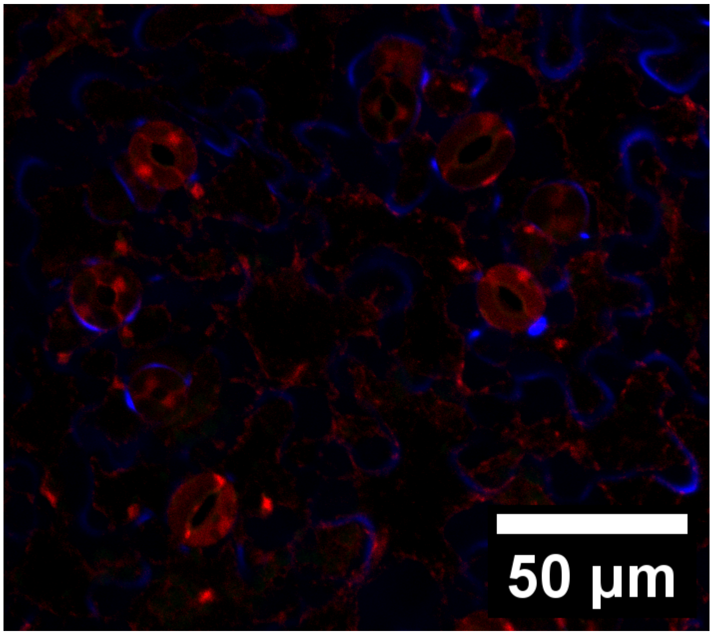


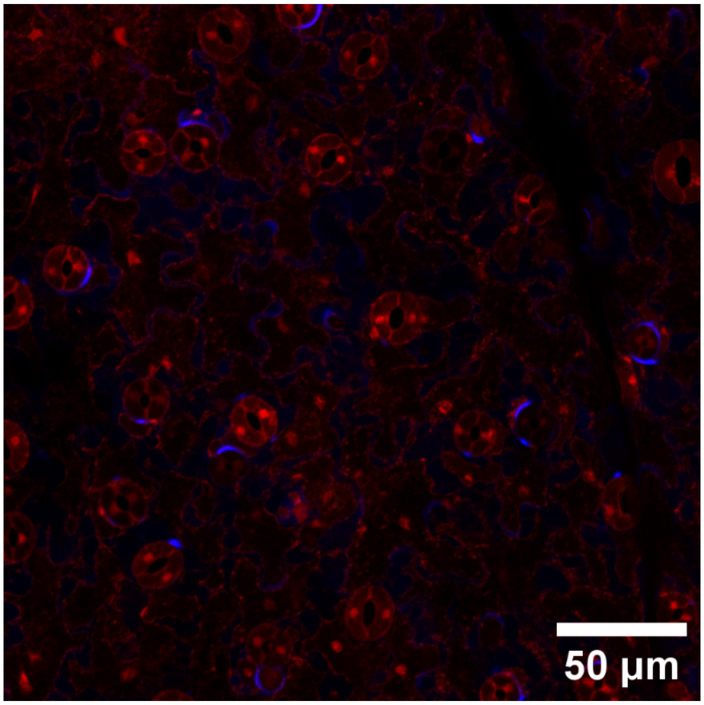

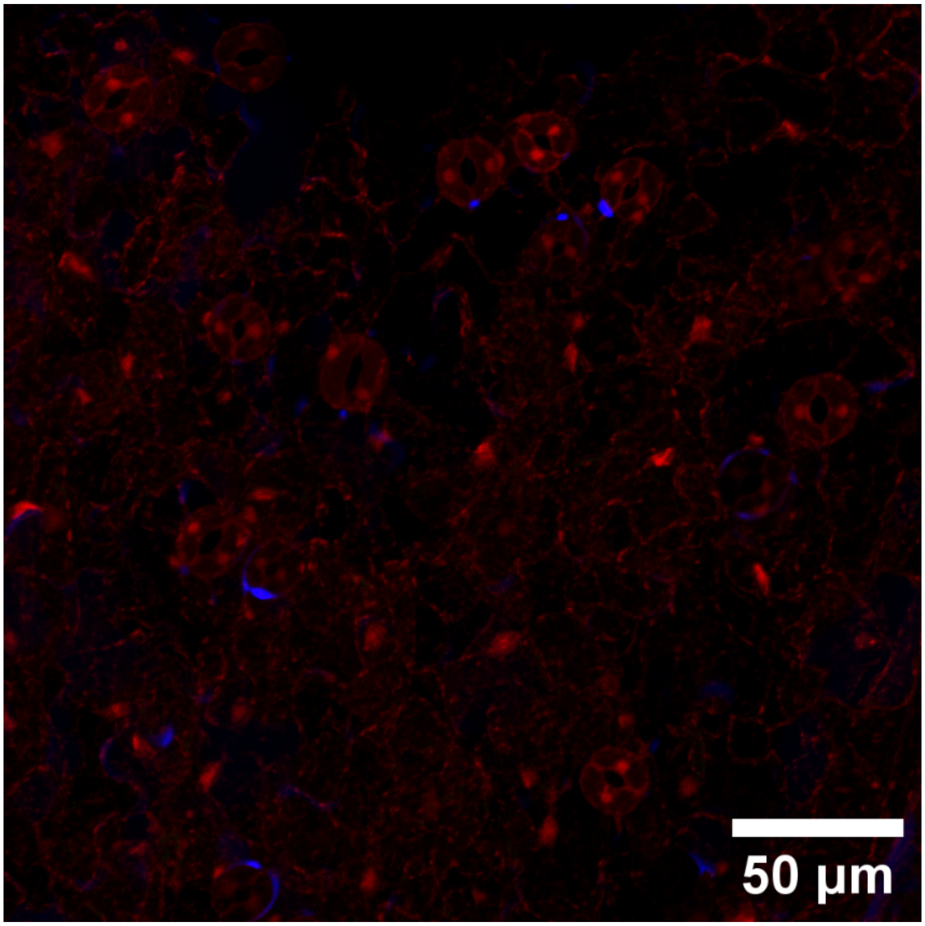


**underswitched**

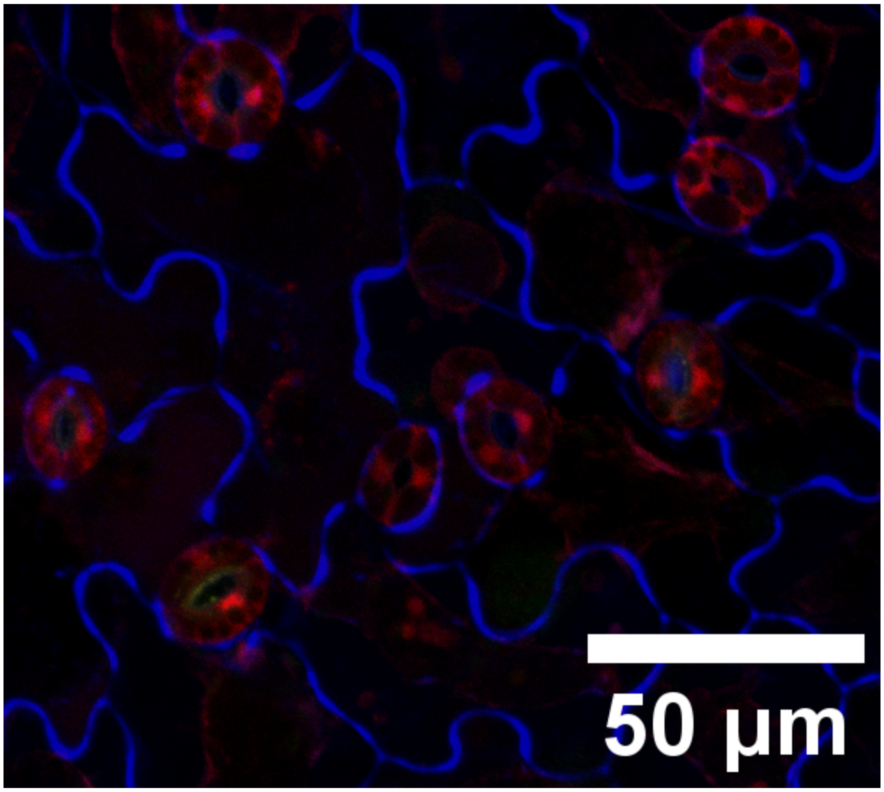

**pMUTE single integrase switch (Bxb1 then PhiC31 target transformed with pMUTE::Bxb1)**

**‘as expected’**

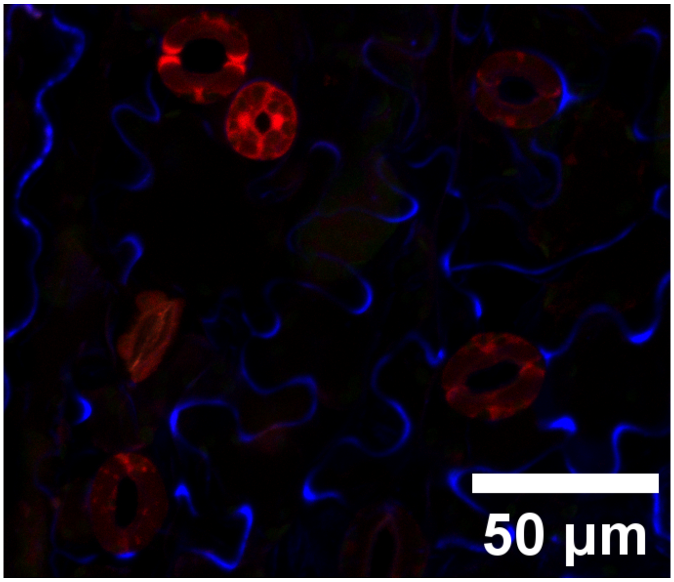


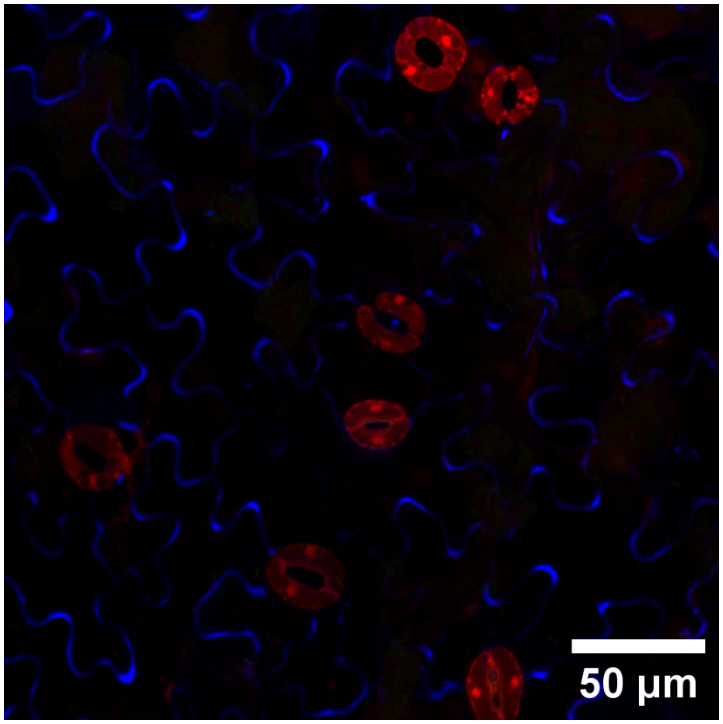


**‘overswitched’**

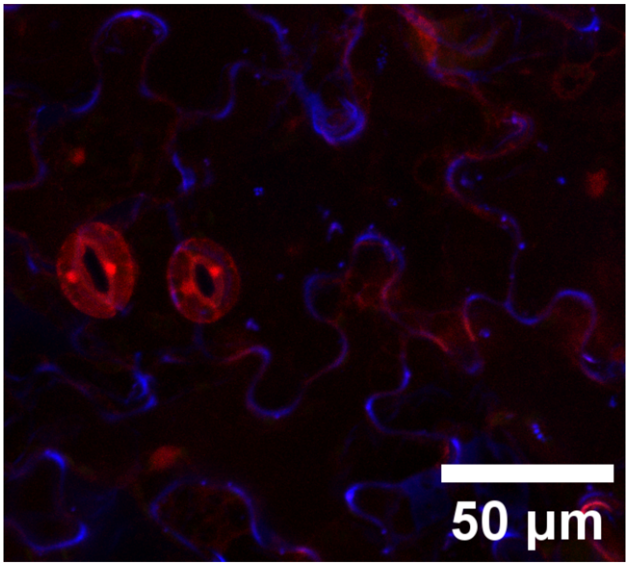


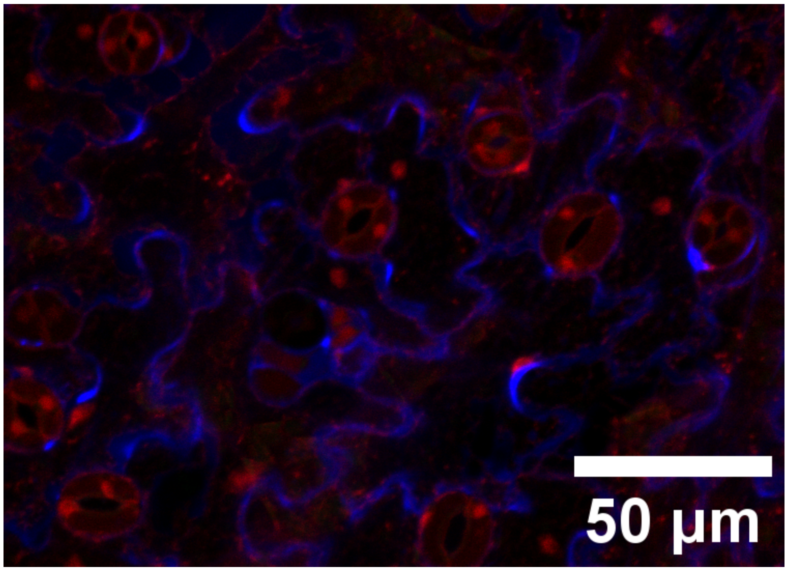

### Full stomata recorder (pSPCH::PhiC31, pMUTE::Bxb1 transformed into the PhiC31 then Bxb1 target)

**'as expected'**

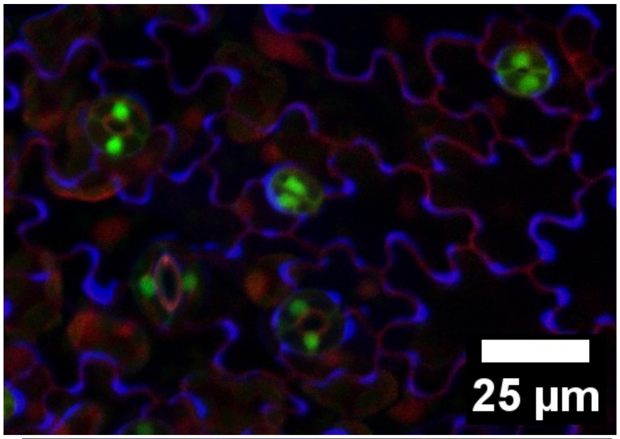


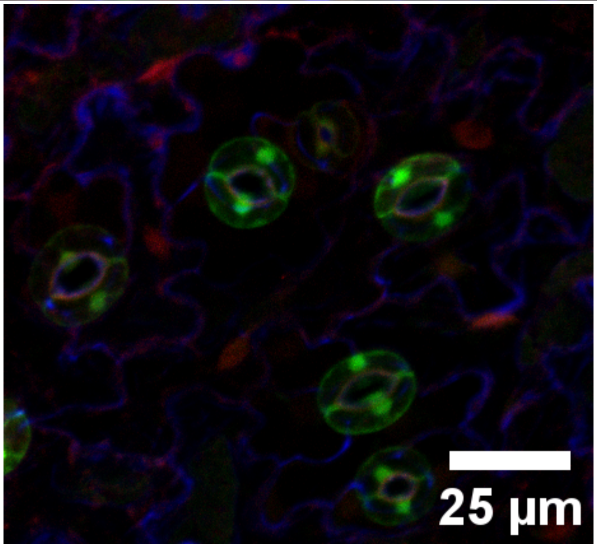


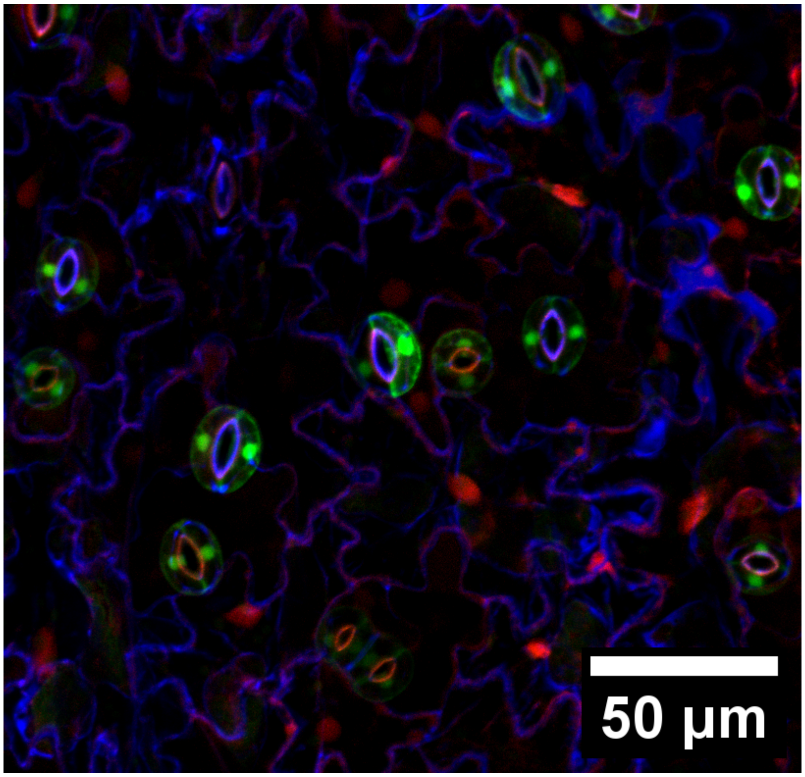


**PhiC31 overswitched**

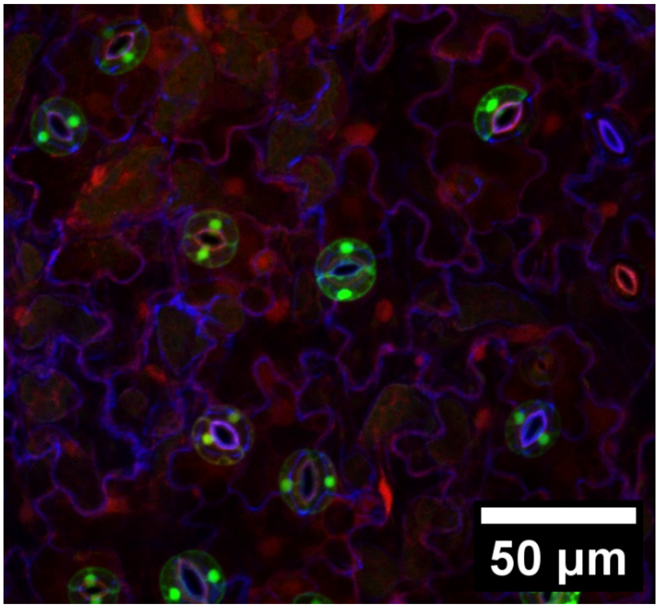

**Bxb1 overswitched**

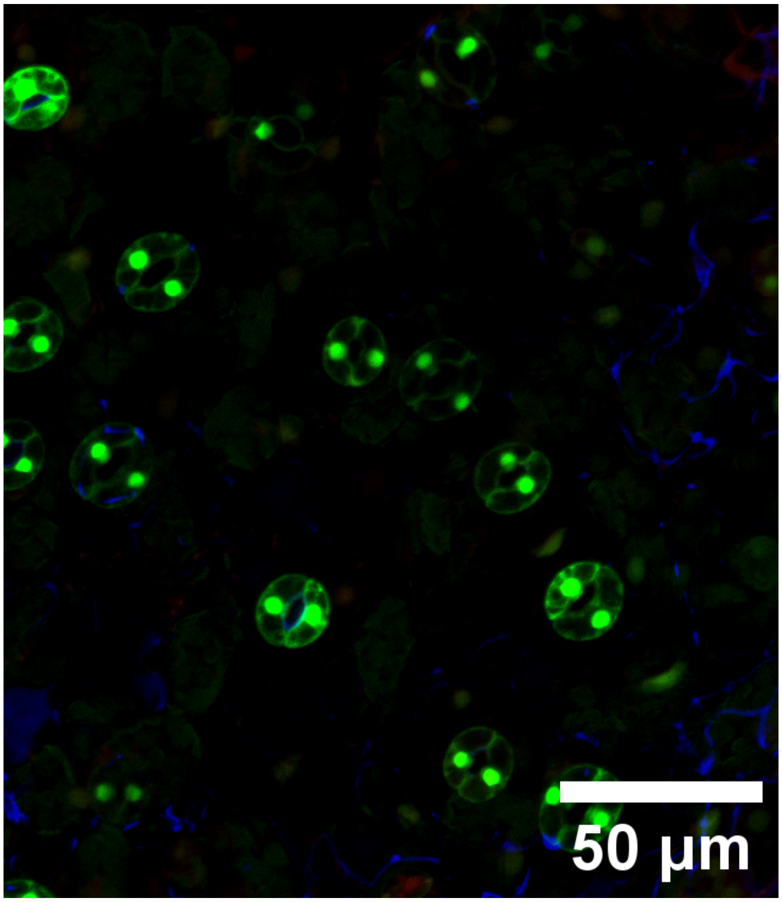


**PhiC31 underswitched**

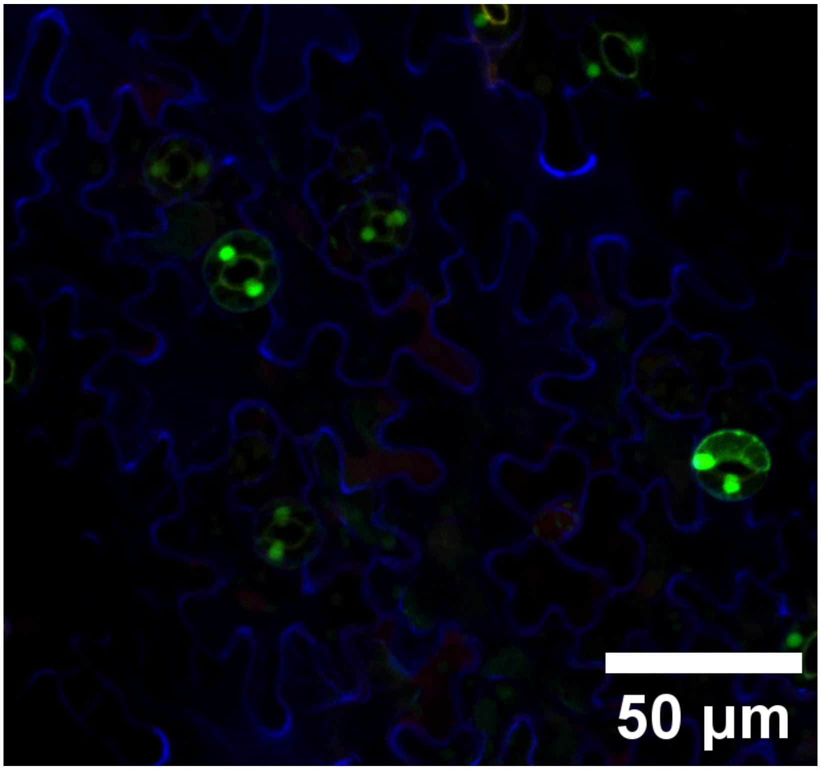
